# Dynamics of age-related catastrophic mitotic failures and recovery in yeast

**DOI:** 10.1101/466797

**Authors:** Matthew M. Crane, Adam E. Russell, Brent J. Schafer, Mung Gi Hong, Joslyn E. Goings, Kenneth L. Chen, Ben W. Blue, Matt Kaeberlein

**Affiliations:** Department of Pathology, University of Washington, Seattle, WA, USA; Department of Genome Sciences, University of Washington, Seattle, Washington, USA; Medical Scientist Training Program, University of Washington, Seattle, Washington USA

## Abstract

Genome instability is a hallmark of aging and contributes to age-related disorders such as progeria, cancer, and Alzheimer’s disease. In particular, nuclear quality control mechanisms and cell cycle checkpoints have generally been studied in young cells and animals where they function optimally, and where genomic instability is low. Here, we use single cell imaging to study the consequences of increased genomic instability during aging, and identify striking age-associated genome missegregation events. During these events the majority of mother cell chromatin, and often both spindle poles, are mistakenly sent to the daughter cell. This breakdown in mitotic fidelity is accompanied by a transient cell cycle arrest that can persist for many hours, as cells engage a retrograde transport mechanism to return chromosomes to the mother cell. The repetitive ribosomal DNA (rDNA) has been previously identified as being highly vulnerable to age-related replication stress and genomic instability, and we present several lines of evidence supporting a model whereby expansion of rDNA during aging results in nucleolar breakdown and competition for limited nucleosomes, thereby increasing risk of catastrophic genome missegregation.

## Main Text

Each cell cycle involves a delicate choreography of duplicating genetic material and cellular organelles, and apportioning them appropriately between mother and daughter cell. Failures of cell cycle regulation can result in severely compromised fitness or cells that respond improperly to environmental cues and emerge as cancerous precursors (Hanahan and Weinberg, 2011). In particular, aneuploidy (the gain or loss of partial or whole chromosomes) can be deleterious to fitness(Beach et al., 2017; Sunshine et al., 2016) and has been implicated in many different types of cancers(Gordon et al., 2012) as well as developmental diseases such as Down Syndrome(Nagaoka et al., 2012). Recent work has also documented extensive damage and genomic rearrangements that can occur from micronuclei or from telomeric crisis(Maciejowski et al., 2015; Zhang et al., 2015), and identified the rDNA sequences as particularly vulnerable to genomic damage(Flach et al., 2014; Xu et al., 2017). Mitotic processes and checkpoints of the budding yeast *Saccharomyces cerevisiae* have been extensively studied, but the vast majority of work has focused on logarithmically growing young cells(Beach et al., 2017; Dotiwala et al., 2007; London and Biggins, 2014; Palmer et al., 1989; Santaguida and Amon, 2015). By studying nuclear dynamics in yeast as they age, we have uncovered a cause of age-associated genomic instability and an active mechanism to maintain nuclear integrity and proper segregation of genetic material in aged cells.

We characterized the dynamics of genome replication and partitioning during replicative aging by imaging cells expressing fluorescently tagged histone 2B (Htb2:mCherry) in a microfluidic device over their entire lifespans(Crane et al., 2014). During each cell cycle, the amount of Htb2 in the mother cell nucleus increases by about two-fold and then drops by one-half, as the cell enters mitosis and chromosomes are segregated to the newly formed daughter. The vast majority of cell divisions in young cells follow this characteristic pattern (Figure 1A). As cells age, however, abnormal segregation events become common (Figure 1B, Videos 1-2, please ensure volume is on for all Videos to hear audio explanation). The single cell trace shown in Figure1C, for example, shows a cell undergoing multiple cell cycles with proper division until an abnormal segregation occurs in which the majority of detectable histones are sent to the daughter cell. These genome-level missegregation (GLM) events result in cell cycle arrest that can range from a few minutes (Figure 1C-top) to many hours (Figure1C-middle), before they are usually corrected by returning the aberrantly segregated genetic material to the mother cell. The range of arrest durations is broad, with most events resolved within an hour, but some lasting many hours (Figure 1 – figure supplement 1). If corrected by this REtrograde TRansport Nuclear (RETRN) process, mother cells are able to proceed through subsequent divisions, but if not, the mother cells will terminally exit the cell cycle and senesce (Figure 1C-bottom).

**Fig. 1.**
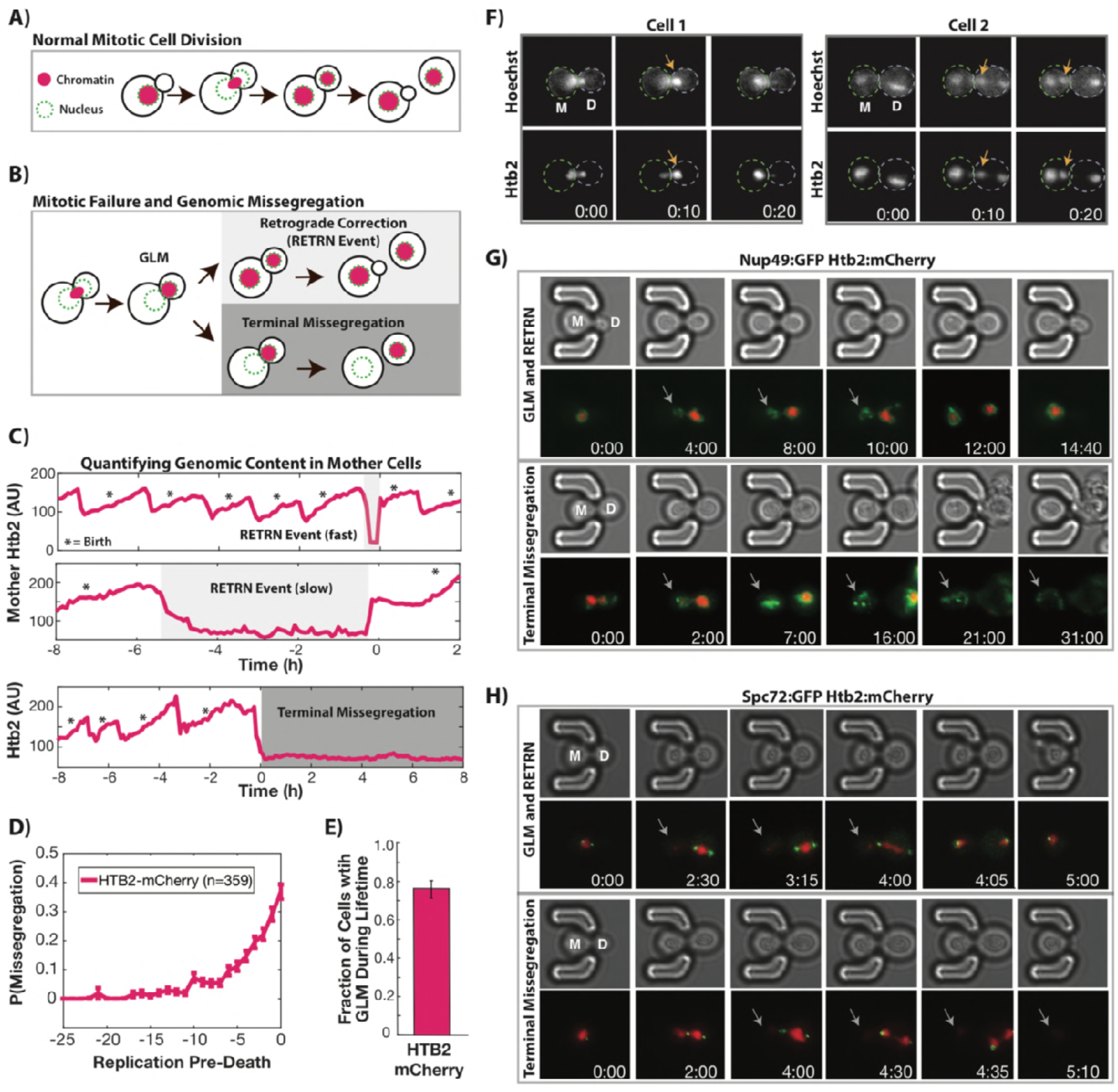
During replicative aging cells frequently undergo dramatic genomic missegregation events. **A**) Schematic showing the process of a normal cell division where chromatin (red) doubles during S-phase and is divided between mother and daughter during mitosis. **B**) Aging cells frequently experience Genome Level Missegregation (GLM) events where most genomic material enters the daughter while the nuclear envelope stays behind. Usually this missegregetion is corrected (top, RETRN event), allowing mother cells to go on to divide and produce more daughters. If not corrected and cytokinesis occurs (bottom), this becomes a terminal event wherein mother cells replicatively senesce. **C**) Representative single cell traces of mother Htb2 levels showing missegregation (shaded) and active retrograde correction events. Corrections can occur quickly (top), or can take hours to be completed (middle). A GLM becomes terminal (bottom) if it is not corrected. (*) indicates the formation of new buds, and both cells with RETRN events produce additional daughters. AU indicates arbitrary units. **D**) Missegregation probabilities increase dramatically near the end of replicative lifespan. n=359 mother cells examined. **E**) Over their entire replicative lifespan, individual mother cells have a greater than 70% chance of having one or more missegregation events. **F**) Genomic DNA and histones co-localize during GLM events. Two cells expressing Htb2:mCherry and stained with a live DNA dye Hoechst 3342. **G**) Time-lapse dynamics of a GLM with RETRN correction (top, mother cell replicative age 14) and a terminal missegregation (bottom, mother cell replicative age 12) in cells co-expressing Htb2:mCherry and Nup49:GFP. During both GLMs the nuclear envelope is clearly visible in both mother (M) and daughter (D) cells. See Videos 6 and 7. **H**) Time-lapse dynamics of a GLM with RETRN correction (top, mother cell replicative age 13) and a terminal missegregation (bottom, mother cell replicative age 16) in cells expressing Htb2:mCherry and Spc72:GFP. Both spindle poles can be seen to enter the daughter (D) during these events, and during the RETRN event a spindle pole returns to the mother (M). In the terminal missegregation, the spindle pole fails to reenter the mother cell. See Videos 8 and 9. Times are indicated in hours:mins from the start of the displayed time-lapse, not the start of the experiment. Arrows indicate mother cells without visible chromatin.

To further characterize GLM and RETRN dynamics during aging, cells were imaged over their entire replicative lifespan, with birth events, GLMs, and RETRN events assessed. The probability of a GLM increased dramatically at the end of life (Figure 1D), with approximately three quarters of mother cells experiencing one or more GLMs (Figure 1E). About 90% of GLMs were corrected through successful RETRN events, allowing individual mother cells to live approximately 30% longer on average than if all GLMs were terminal (Figure 1 – figure supplement 2). However, even when corrected, mother cells that undergo a GLM are more likely to die in the near future than cells of the same age that have not experienced such an event, and GLMs become increasingly predictive of impending mortality with increasing age (Figure 1 – figure supplement 2). To confirm that the histones do indeed co-localize with DNA during these events, we imaged old mothers and observed the dynamics of Htb2 in cells exposed to the DNA stain Hoechst 3342. As can be clearly seen (Figure 1F, Videos 3-4), both the DNA and histones move in concert during these events. Furthermore, we confirmed that histone levels reflect nuclear DNA abundance in single cells by staining with DAPI and comparing Htb2 levels with DNA content in old cells (Figure 1 – figure supplement 3). GLM and RETRN dynamics were not influenced by the fluorophore used or which histone is tagged, as the dynamics of both Htb2:mCherry and histone 2A tagged with GFP (Hta2:GFP) did not differ (Figure 1 – figure supplement 4). GLM and RETRN frequency is not an artifact of our imaging protocol, as modifying the excitation power or the cumulative excitation energy exposure had no effect on these observations (Figure 1 – figure supplement 5). For clarity, the strain containing Htb2:mCherry is referred to as wild-type hereafter.

In order to observe the nuclear periphery during GLM and RETRN events, we imaged aging cells expressing both Htb2:mCherry and Nup49:GFP, and compared normal divisions (Video 5) with RETRN events (Figure 1G-top, Video 6) and terminal GLMs (Figure 1G-bottom, Video 7). The dynamics of the histone missegregation and recovery can be clearly seen in these time-lapse series, and strikingly the mother cells retain an intact nuclear envelope during these events – even if they lose all the chromatin (Figure1G). Passage of the histones fully into the daughter cell is evident from cells co-expressing a bud neck marker (Myo1:GFP) along with Htb2:mCherry (Videos 1-2). Interestingly, during these events both spindle poles often enter the daughter and move in concert with the tagged histones (Figure1H). Spindle missegregation and RETRN was visualized through time-lapse imaging of aging cells expressing the spindle component Spc72 tagged with GFP (Spc72:GFP, Video 8). In terminal GLMs without RETRN, both spindle poles generally remain in the daughter cell (Figure 1H, Video 9). This can also be observed in videos where tubulin is tagged with GFP (Tub1:GFP), and all of the detectable nuclear microtubules enter the daughter cell during GLMs (Video 10-11).

To investigate the temporal and cell cycle dynamics of the genome missegregation and RETRN events, we employed two cell cycle reporters, Whi5:GFP and Myo1:GFP(Di Talia et al., 2007). Whi5 prevents exit from G1 when localized in the nucleus, and the transition of Whi5 from the nucleus to the cytoplasm signifies the end of G1 (Figure 2A) (Charvin et al., 2010). Individual young cells proceed in a reliable fashion through the cell cycle, with Whi5 becoming nuclear localized immediately after Htb2 levels fall and the cell enters telophase (FigureS6). By aligning all annotated cell divisions without GLMs (Figure 2B), the temporal dynamics of histone separation into the daughter cells are clear and immediately followed by the transition of Whi5 from the cytoplasm into the nucleus. In contrast, when a GLM occurs, Htb2 levels fall precipitously in mother cells, but the cell delays the cytoplasmic to nuclear transition of Whi5 until after the RETRN correction (Figure 2C). Because the Whi5 cytoplasm-to-nuclear transition occurs upon activation of the mitotic exit network(Bean et al., 2006; Costanzo et al., 2004), this delay demonstrates that the cell prolongs mitosis until the GLM is corrected.

**Fig. 2.**
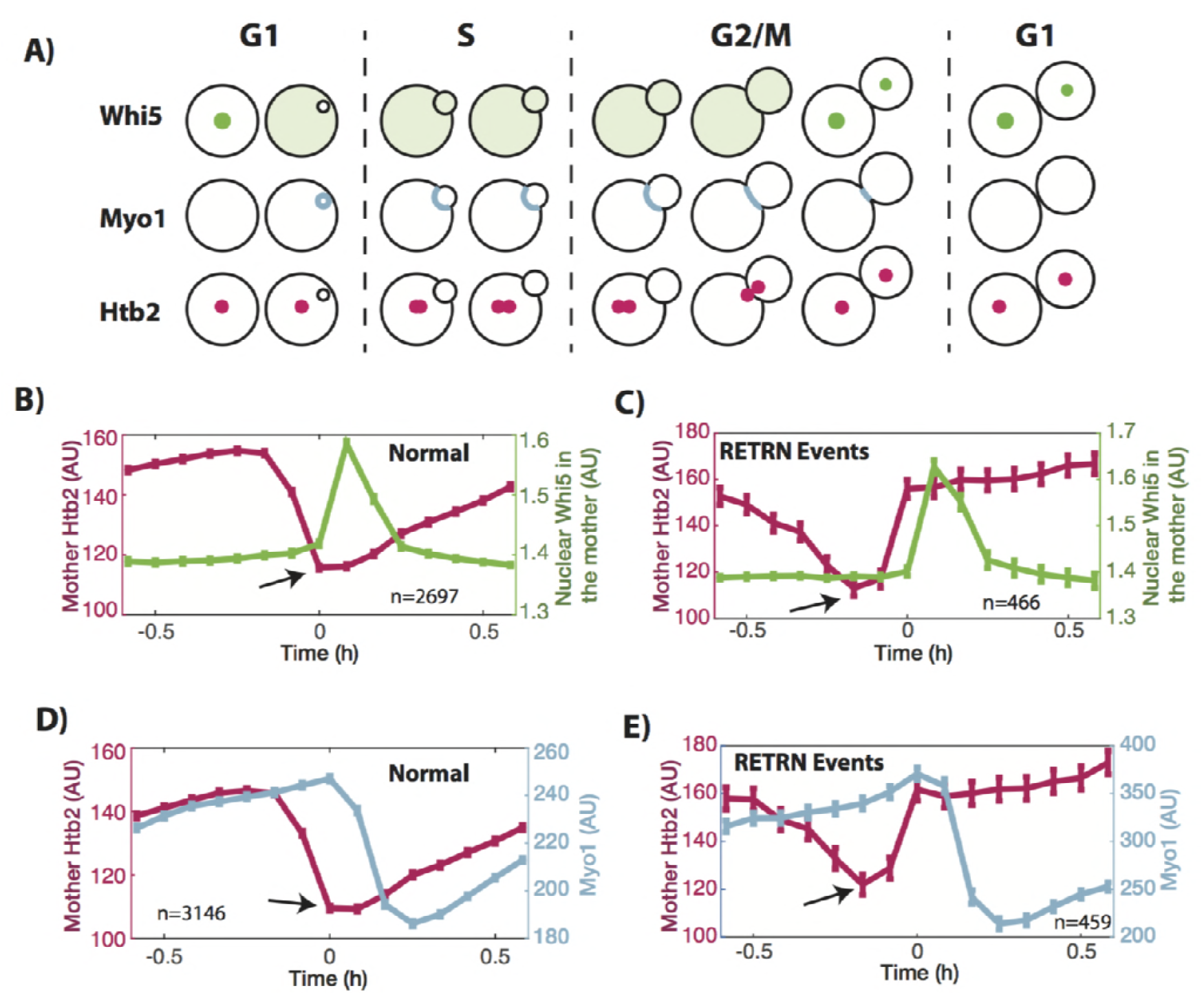
Characterization of missegregation and corrections during the cell cycle. **A**) Schematic showing the temporal dynamics of the proteins used to characterize missegregation events. Whi5 exits the nucleus to initiate START and move the cell into S phase. It re-enters the nucleus at the end of mitosis. Myo1 is produced at the end of G1 and localizes to the bud neck until cytokinesis ends mitosis. **B**) Average Htb2 and Whi5 dynamics for cell cycles without missegregation events (n=2,697). Whi5 begins to transition from cytoplasmic to nuclear when Htb2 levels reach the lowest point as indicated by the arrow. **C**) Average traces from cell cycles with RETRN events. In these cases (n=466), the Whi5 cytoplasmic to nuclear transition is delayed until the missegregation is corrected. **D**) Average Htb2 and Myo1 dynamics for cell cycles without missegregations (n=3,146). Myo1 levels begin to fall immediately after chromosome segregation, as shown by the arrow. **E**) In cell cycles with a RETRN event (n=459), even after chromosomes enter the daughter (noted by the arrow), Myo1 levels continue to increase until the missegregation is corrected, and then cytokinesis begins. N-values indicate the number of cell cycles analyzed, and all error bars are standard error. Cells are aligned so that time t=0 indicates the beginning of mitotic exit and the M to G1 transition, as determined by Whi5 and Myo1 dynamics.

In a complementary fashion, we confirmed that GLMs induced a delay in mitotic exit by following Myo1:GFP localization and abundance. Myo1 is produced at the end of G1 as the cell begins the formation of a new daughter(Bi and Park, 2012; Weiss, 2012). At the end of mitosis, following successful partitioning of the genomic material, Myo1 is degraded during cytokinesis (Figure 2A, Figure 2 – figure supplement 1). By manually aligning normal cell divisions, the increase of Myo1 levels at the bud neck, and subsequent decrease during cytokinesis, can be clearly seen (Figure 2D). When the Htb2 levels reach their lowest point, Myo1 levels are at a maximum, but they immediately begin to fall as the cell eliminates the bud neck during cytokinesis. In contrast, in divisions where a GLM occurs, the levels of Myo1 continue to increase even after the Htb2 levels reach their lowest point (Figure 2E). Only following the RETRN event does Myo1 begin to fall.

To determine whether GLMs result from improper spindle attachment and whether RETRN events are caused by activation of the spindle assembly checkpoint, we deleted the gene encoding the spindle assembly checkpoint component Mad3 (mammalian BubR1). This failed to alter the age-related increase in missegregation, and older *mad3*Δ cells had the same missegregation rates as wild type cells (Figure 2 – figure supplement 2). Taken together, these data differentiate RETRN events and terminal GLMs from prior observations of nuclear oscillations near the bud neck linked to alignment of the spindle poles between mother and daughter(Palmer et al., 1989; Yang et al., 1997; Yeh et al., 2000, 1995). Similarly, nuclear excursions where a single spindle-pole entered the daughter, but then returned to the mother has been identified in DNA damage checkpoint mutants (Dotiwala et al., 2007); however, when cells arrest as a result of the DNA damage response during metaphase, only the nucleolus enters the daughter while all chromatin is retained by the mother cell (Witkin et al., 2012). A recent report identified segregation of the nucleus and spindle poles into the daughter cell in five aging yeast cells (of the 10 observed), which is likely to be the same phenotype detailed here(Neurohr et al., 2018). They also identified elevated rates of chromosome I missegregation in old cells (in ten of the forty cells observed). In prior studies, mutants that missegregate chromosomes or both spindle poles to the daughter cell are unable to correct these events(Finley et al., 2008; Thrower et al., 2003; Yeh et al., 2000, 1995); however, the RETRN events seen in aged mother cells unambiguously delay mitotic exit and correct these failures by actively transferring chromatin, microtubules, and/or spindle poles from the daughter cell to the mother cell.

Changes in nucleosome occupancy have been linked to age-related genomic instability(Hu et al., 2014), DNA damage(Hauer et al., 2017), and tumorigenesis(Sharma et al., 2010), and during the course of our studies we observed increasing Htb2 levels at late replicative ages (Figure 3A, Video 12). Given that prior studies have reported both increasing and decreasing histone levels with age(Feser et al., 2010; Janssens et al., 2015), we analyzed expression of two histones tagged with GFP (Hta2, Htb2) in addition to Htb2:mCherry and confirmed that this end of life increase is a general behavior of histones (Figure 3 – figure supplement 1). Furthermore, all histones examined showed a dramatic increase in expression between 5-10 divisions prior to death (Figure 3 – figure supplement 2).

**Fig. 3.**
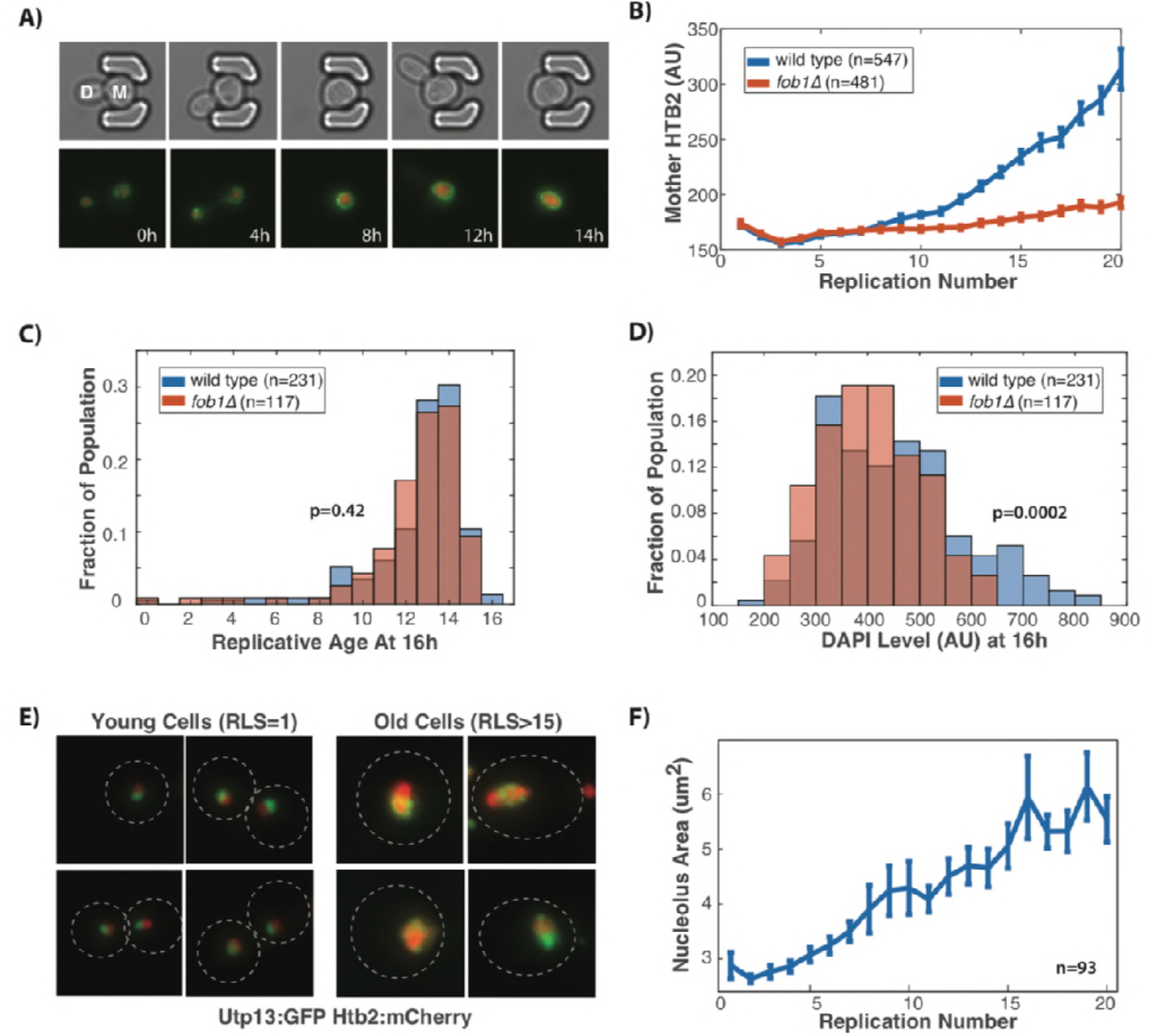
Age-related rDNA instability impairs Cdc14 release and anaphase entry. **A**) Time course of a single mother cell expressing Htb2:mCherry and Nup49:GFP shows the increase in histone levels during normal replicative aging, full movie in Video 12. The mother cell shown here has a replicative age of 16 at the timelapse start and 21 at the end. **B**) Old wild type cells have significantly greater histone levels compared to young wild type cells and accumulation of histones during aging is attenuated in cells lacking Fob1 (n=481) compared to wild type (wt, n=587). Significance of age and genotype (p<0.0001 in both cases) determined by repeated measures ANOVA. This links rDNA expansion in the form of ERCs with histone accumulation. **C**) Wild-type and *fob1*Δ cells were grown in the microfluidic device for 16h, and then fixed. At the time of fixation, both strains had similar replicative ages (p=0.42, two tailed t-test). **D**) Although both strains had similar replicative ages, the aged wild-type mother cells had increased DAPI staining levels relative to aged *fob1Δ* cells, indicating an increased level of DNA (p=0.0002, two tailed t-test). **E**) In young cells the nucleolus (Utp13:GFP) is adjacent to the genomic DNA (marked by Htb2). In aged cells, however, ERCs accumulate and are localized to the nucleolus which becomes colocalized with histones as the nucleolus fragments. Colocalization is particularly evident toward the end of a mother cells life (see Videos 13 and 14). **F**) The size of the nucleolus as measured by segmenting Utp13:GFP increases during replicative aging (n=93, p<0.0001, Mann-Kendall trend test).

We speculated that the elevated histones in aged cells may reflect an expansion of the rDNA in the form of extrachromosomal rDNA circles (ERCs), which increase dramatically during aging and have been proposed as a molecular mechanism of aging(Sinclair and Guarente, 1997) and genome instability (Ganley et al., 2009). As a result, rDNA copy number in old cells has been shown to increase nearly twenty fold (Dang et al., 2009). To test whether histone levels reflect ERC abundance, we determined the impact of reducing ERC formation by removing the replication fork block protein Fob1; deletion of *FOB1* reduces rDNA recombination and ERC abundance(Defossez et al., 1999; Kobayashi et al., 1998; Sinclair and Guarente, 1997) and significantly extends lifespan (Figure 3 – figure supplement 3). In comparison to wild type cells, the increase in histone levels with age is attenuated in *fob1*Δ cells (Figure 3B). Furthermore, although wild-type and *fob1*Δ mother cells stained with DAPI after 16h of growth had statistically indistinguishable replicative ages (Figure 3C, median RLS=13, p=0.42), wild-type cells have significantly greater DNA levels (Figure 3D, p=0.0002). The lower DNA and histone levels in old *fob1*Δ cells relative to age-matched wild-type is consistent with the model that increasing histone levels are driven by ERC accumulation in aging yeast. If increasing histone levels reflect ERC levels, then histone levels should act as a biomarker that predicts remaining lifespan and could be associated with other age-associated failures. Indeed, within cells of the same replicative age, we determined that histone abundance is negatively correlated with remaining lifespan, and this correlation becomes stronger as cells age (Figure 3 – figure supplement 3). Histone levels are less correlated with mortality in *fob1*Δ cells, supporting a model where slowing ERC production reduces the underlying pathology linking histone abundance to death.

The rDNA is localized to the nucleolus, a phase separated organelle inside the nucleus(Lindström et al., 2018). To examine the relationship between histone dynamics, ERCs, and nucleolar structure, we followed aging mother cells expressing Htb2:mCherry along with a GFP fusion to the nucleolar protein Utp13. In young cells, the nucleolus appears as a nuclear site adjacent to the majority of histones, while in old cells, a large quantity of histones is localized within the nucleolus (Figure 3C, Videos 13-14). Furthermore, the size of the nucleolus increases dramatically as cells age and often becomes fragmented into multiple foci (Figure 3D, Videos 13-14), further indicating that the excess histones are localized to ERCs in old cells and confirming earlier work linking ERCs to nucleolar fragmentation(Neurohr et al., 2018; Sinclair et al., 1997; Sinclair and Guarente, 1997).

The protein Cdc14 is also localized to the nucleolus where it functions in the Regulator of Nucleolar Silencing and Telophase Exit (RENT) complex to silence transcription (Clemente-Blanco et al., 2011; Shou et al., 1999) and ensure proper segregation of chromosomes(D’Amours et al., 2004; Rock and Amon, 2009; Sullivan et al., 2004). Recently Cdc14 was identified as the limiting step in anaphase, and separately it was observed that compaction of rDNA within the nucleolus interfered with proper release of Cdc14 from the nucleolus(de los Santos-Velázquez et al., 2017; Roccuzzo et al., 2015). Given the increased nucleolar size and the increased nucleolar histone levels during aging, we hypothesized that this might affect proper Cdc14 release from the nucleolus during anaphase. To test this, we observed both Cdc14:GFP and Htb2:mCherry dynamics in individual cells, and compared normal cell cycles with cell cycles where GLMs occurred. Cdc14 exit from the nucleolus is controlled in two stages, first by the CDC Fourteen Early Anaphase Release (FEAR) network and then the Mitotic Exit Network (MEN)(Rock and Amon, 2009). By averaging the dynamics of cell cycles that behave normally, it is clear that Cdc14 begins to exit the nucleolus prior to division of genomic material between mother and daughter cells (Figure 4A, Video 15). In cell cycles that experience GLMs, however, Cdc14 remains localized to the nucleolus but is released immediately preceding a RETRN event (Figure 4B, Video 15). This agrees with prior work showing that Cdc14 release during anaphase is required to generate pulling forces within the mother to counteract those in the daughter (Ross and Cohen-Fix, 2004).

**Figure 4.**
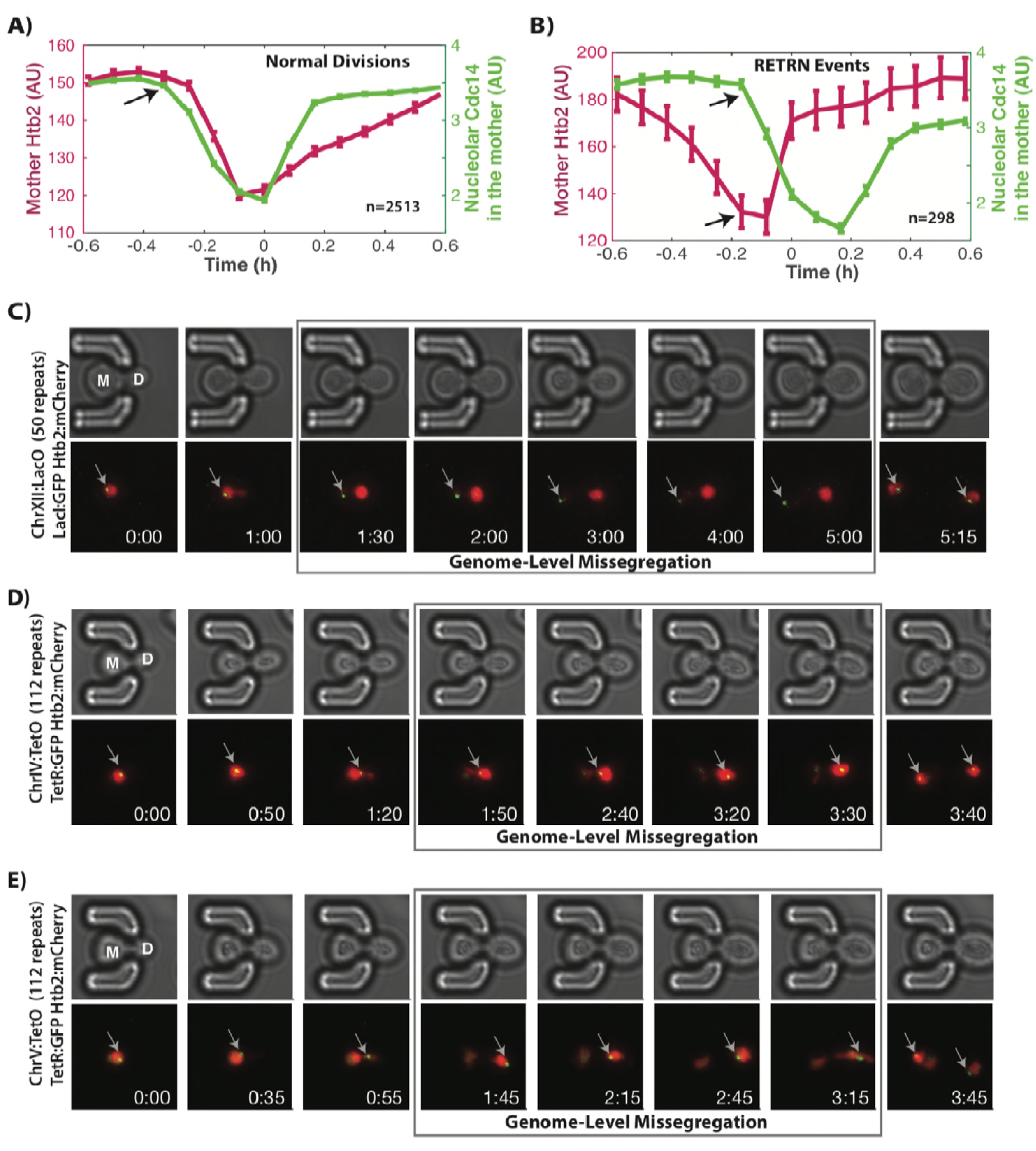
Altered Cdc14 dynamics during aging disrupts segregation of rDNA containing Chromosome XII. **A**) Dynamics of Cdc14 and Htb2 in mother cells during normal divisions (n=2513). As the cell enters mitosis, Cdc14 leaves the nucleolus which triggers anaphase (indicated by the arrow). **B**) Dynamics of Cdc14 and Htb2 in mother cells during RETRN events (n=298). Although genomic content has left the mother and entered the daughter, Cdc14 is still localized to the nucleolus. Shortly following Cdc14 release from the nucleolus (indicated by the arrows), the cells experience a RETRN event. **C**) Direct observation of Chr XII using LacI:GFP and LacO repeats on Chr XII. When the cell experiences a missegregation event, both chromatids of Chr XII remain behind in the mother until the RETRN event. After this, a single green dot can be seen in both mother and daughter cells. Mother cell replicative age equals 8 at the beginning of this timelapse. **D**) Direct observation of Chr IV using TetR:GFP and TetO repeats on Chr IV. When the cell experiences a GLM, both chromatids of Chr IV move to the daughter along with the majority of the chromatin. Following a RETRN event, single green dot can be seen in both mother and daughter cells. Mother cell replicative age equals 5 at the beginning of this timelapse. **E**) Direct observation of Chr V using TetR:GFP and TetO repeats on Chr V. When the cell experiences a GLM, both chromatids of Chr V move to the daughter along with the majority of the chromatin. Following a RETRN event, single green dot can be seen in both mother and daughter cells. Mother cell replicative age equals 19 at the beginning of this timelapse. The gray arrows mark the location of the labelled chromosomes. Times are indicated in hours:mins. In all panels, n-values report the number of individual cells analyzed.

Cdc14 is specifically required for condensation and segregation of repetitive DNA sequences including the rDNA and telomeres(D’Amours et al., 2004; Sullivan et al., 2004). In order to further explore the consequences of failed Cdc14 release on anaphase dynamics during GLM events, we directly observed Chromosome XII by targeting a LacI:GFP reporter to LacO sites engineered on Chr XII(Ide et al., 2010). During GLMs where the majority of DNA enters the daughter cell, both Chr XII chromatids remain behind in mother cell (Figure 4C, Video 16-17). Furthermore, during these GLMs, the Chr XII sister chromatids appear as a single point, only separating into two distinct foci following a RETRN event (Figure 4C, Video 16-17). This suggests that delayed Cdc14 activation prevents separation of the rDNA and results in improper condensation and segregation of Chr XII. During this process, as Chr XII remains localized to the nucleolus in the mother cell, the remaining genomic content missegregates to the daughter. We confirmed that this behavior is unique to Chr XII by imaging Chr IV and V during GLMs. Both copies of Chr IV and Chr V are missegregated to the daughter cells with the rest of the chromatin during GLMs (Figure 4D,E, Videos 18-19). Thus, we propose that age-associated expansion of the rDNA and histone depletion lead to dysregulation of Cdc14 during the metaphase to anaphase transition which results in improper genomic segregation. In both terminal GLMs and RETRN events, Cdc14 eventually exits the nucleolus to trigger anaphase (Figure 4 – figure supplement 1).

During mitosis, cells must remove and then replace nucleosomes on each DNA strand. Because all replicating DNA pulls histones from a common pool, a mismatch between local histone supply and demand could lead to gaps in nucleosome occupancy, which have been previously observed in aged cells(Hu et al., 2014). We hypothesized that expansion of the rDNA due to increasing ERC levels during aging elevates mother cell demand for histones, which is only partially compensated for by the observed increase in histone expression. By this model, histone demand should be lower in old *fob1*Δ cells compared to age-matched wild-type cells (due to fewer ERCs), which should result in a reduced probability of GLMs. Observations of GLMs in a *fob1*Δ mutant support this model (Figure 5A). In particular, *fob1*Δ and wild-type cells diverge most dramatically following replicative age 10 (Figure 5A), when the wild-type cells begin to increase histone levels (Figure 3B).

**Figure 5.**
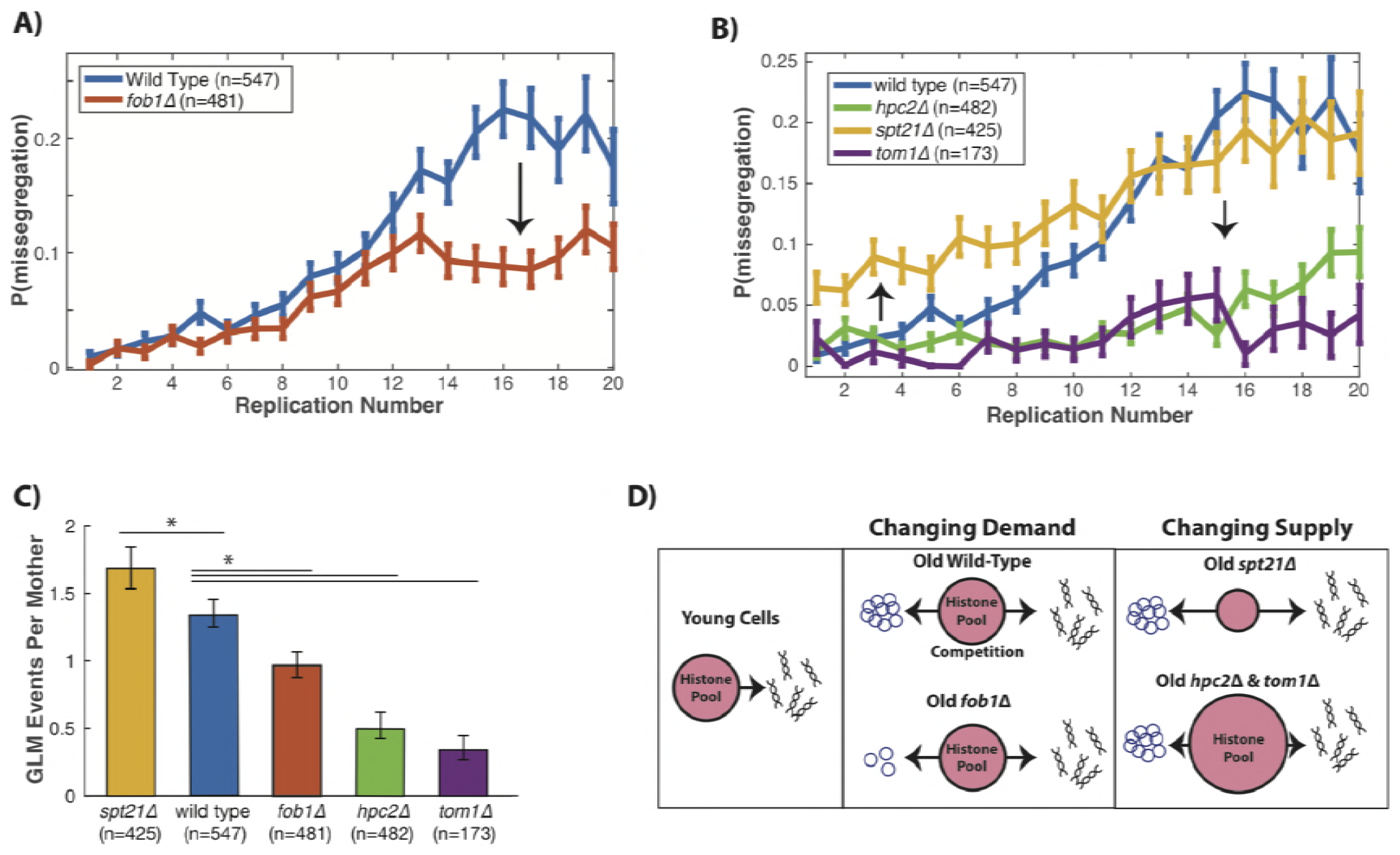
Genomic missegregations increase with age as a result of competition for histones. **A**) Deleting *FOB1* reduces the accumulation of ERCs with age, and also reduces the probability that cells will experience a missegregation event, shown as P(missegregation) at different ages. **B**) Increasing histone supply (*hpc2*Δ & *tom1*Δ) reduces the fraction of cells that have missegregation events, while reducing histone supply (*spt21*Δ) increases the fraction of cells with missegregation events. Error bars are standard error. N-values report the number of cells. **C**) The mean number of genome level missegregations (GLM) events per mother cell by replicative age twenty. Error bars are 95% confidence intervals generated by bootstrapping with replacement. (*) Indicates samples with confidence intervals that do not overlap with wild-type. **D**) In young cells the common pool of histones is available for genomic DNA, but as cells age the demand for histones increases as ERCs compete with genomic DNA for the free pool of histones. Old *fob1*Δ cells have fewer ERCs which lowers the competition for histones. By increasing the common histone supply (*hpc2*Δ and *tom1*Δ) competition between genomic DNA and ERCs is reduced and the missegregation rate is reduced.

To directly test the mechanistic link between histone competition and GLMs, we genetically manipulated the supply of histones (Figure 5B, Figure 5 – figure supplement 1). To increase histone levels, we first removed Hpc2, a component of the HIR complex which represses histone transcription(Green et al., 2005). Deletion of *HPC2* results in reduced frequency of GLMs in aging cells (Figure 5B). Likewise, deletion of *TOM1*, which encodes a factor required for degradation of excess histones(Singh et al., 2009), also reduced GLMs (Figure 5B). To reduce histone abundance, we deleted *SPT21*, which encodes a protein that positively regulates histone expression(Dollard et al., 1994; Kurat et al., 2014) and whose loss has been previously shown to reduce histone levels and increase rDNA instability(Eriksson et al., 2012; Kobayashi and Sasaki, 2017). In contrast to deletion of either *HPC2* or *TOM1*, deletion of *SPT21* caused increased frequency of GLMs in aging mother cells. When averaged over the entire lifespan, *spt21Δ* cells experienced significantly more GLMs than wild-type cells, while *hpc2Δ* and *tom1Δ* cells had significantly fewer events (Figure 5C). These observations demonstrate that altering histone abundance in aging cells is sufficient to modulate the frequency of GLM events both upward and downward, and support a model whereby rDNA expansion causes increasing competition for a common histone pool in aging cells which drives the dramatic increase in GLMs during aging (Figure 5D).

Because function declines in many different and subtle ways during aging, catastrophic failures and homeostatic systems like the those uncovered here may only be detected in aged organisms. Imaging of individual yeast cells through microfluidic trapping allowed us to observe GLMs that occur in most mother cells at some point during their lives. These events are exceedingly rare in young cells, are actively repaired through transient arrest of the cell cycle, and likely result from insufficient histone availability due to expansion of the nuclear genome via accumulation of rDNA. The rDNA has been linked to genomic instability in yeast(Ganley et al., 2009; Ide et al., 2010; Saka et al., 2013) and within cancer cell lines(Xu et al., 2017), and nucleolar size is anti-correlated with lifespan across organisms(Tiku et al., 2016). Nucleolar structure has also been linked to cancer pathogenesis(Lindström et al., 2018). Given the ubiquity of aneuploidy in age-related human cancers(Sansregret and Swanton, 2017), we speculate that mechanisms for responding to mitotic failures may be important to cope with age-associated genomic instability in multicellular eukaryotes. Interestingly, although humans have only five acrocentric chromosomes with rDNA repeats, these are far more likely to result in developmental trisomy than other chromosomes(Nagaoka et al., 2012). Whether the mechanisms accounting for age-related increases in genomic instability uncovered here, and the RETRN process to repair these events, are conserved in higher eukaryotes requires further investigation. Such research into the behavior of natural cellular processes during aging could provide insights into age-related pathology and uncover potential targets for intervention.

## Acknowledgements

We would particularly like to thank S. Biggins, B. Brewer and M. Raghuraman for constructive discussions. We also thank L. Veenhoff and Kaeberlein lab members for feedback and advice. Strains YSI129, AMY914 and AMY1081were generous gifts from Jessica Tyler and Adele Marston. This work was supported by NIH grants T32AG000057, R01AG056359, and P30AG013280.

## Supplementary Materials and Methods

### Microfluidics

Cells were imaged using a PDMS microfluidic flow chamber modified from an earlier design (Crane et al., 2014) to increase retention over the replicative lifespan of the mother cells. The microfluidic device was composed of multiple chambers in the same fashion as (Granados et al., 2017), which allowed individual genotypes to be exposed to identical environments and imaged in the same experiment while being physically isolated. Cells were loaded according to previously published methods (Granados et al., 2017). A volumetric flow rate of 3-7 μL/min per chamber was used, with the flow rate starting low, and increasing during the experiment to improve mother cell retention and to ensure that cells do not aggregate, which can clog the device.

### Microscopy

Cells were imaged using a Nikon Ti-2000 microscope with a 40X oil immersion objective with a 1.3 NA and using the Nikon Perfect Focus System. An enclosed incubation chamber was used to maintain a stable 30C environment for the duration of the experiment. Two Aladdin syringe pumps were used for media flow. An LED illumination system (Excelitas 110-LED) was used to provide consistent excitation energies, and to minimize the exposure, illumination was triggered by the camera. Images were acquired using a Hammamatsu Orca Flash 4.0 V2. The microscope was controlled by custom software written in Matlab^®^ and Micromanager.

Images were corrected for illumination artifacts in two stages. First, to correct for individual differences in the pixel biases, 1,000 images were acquired with no illumination, and the individual pixel means were determined. Second, to correct for flatness of field, a fluorescent dye was added to a microfluidic device instead of using a slide with dye. Using a slide containing dye introduces a large amount of out-of-focus light, which results in an underestimation of the field curvature. In order to compensate for the microfluidic features, 1,000 images were acquired each with a small offset in the x and y positions. Images were then dilated, and the median value at each location was used. Thus, for each image, the camera bias for that pixel was subtracted, and then it was multiplied by a flatness of field correction factor.

Images were acquired at 5 min intervals for bright-field and fluorescent channels. The fluorescence excitation power was set to 25% for all imaging except the GFP tagged histones, where it was set to 12%. Fluorescence and brightfield light was activated during image acquisition and all other lights in the room were turned off. For bright-field, 3 z-sections were acquired with 2.5 μm intervals, exposure times of 30 ms and were used for automated segmentation and tracking. For the fluorescent channels, 3 z-sections were acquired with 1.5 μm spacing. GFP images were acquired using a Chroma ET49002 filter set, and mCherry images were acquired using a Chroma ET49306. GFP images were acquired using exposure times of 60ms for all proteins except Htb2 and Hta2 which were acquire using a 30ms exposure time. mCherry images were acquired using a 60 ms exposure time. These imaging conditions were found to work as a reasonable compromise between the desire for frequent, dense imaging to enable identification of missegregations and retrograde transport, while also minimizing phototoxicity. We performed control experiments to verify that these exposure conditions did not affect the rates of genomic missegregation or replicative lifespan (Figure 1 – figure supplement 5). Each strain was imaged in multiple independent experimental runs, each with approximately equal numbers of cells.

### Data processing and single cell scoring

Following data acquisition, cells were identified and tracked using previously published software(Bakker et al., 2017). This identified the cell outline, and performed initial tracking of the cells through time. To ensure that only young, healthy cells were assessed, we only used cells that were identified in the first three hours of the experiment. Birth events for these cells were then manually scored, and any errors in tracking were corrected. This was all done using the bright-field images. Birth events were scored by multiple observers who were blinded. Because individual cells can be lost from traps prior to death, it can be challenging to know whether using censored (lost) cells is most appropriate. In the supplementary information, all data is presented with and without censoring. In the main text, plots aligned based on increasing age used all cells present at that age, even if they were later lost from the device. For plots aligned by death, only cells that had either died or senesced (failed to initiate a new cell division but did not visibly lose cell wall integrity during the experiment) were used. Because censoring in lifespan experiments relies on the assumption that losses are unbiased, we provide replicative lifespan curves both including and excluding censored cells for all strains. Censoring does not change the interpretation or statistical outcome of any of the experiments presented here.

For each cell division, the mean histone levels over that cell cycle were used. For cell cycles that last more than three hours, only the first three hours were used to determine the histone levels. This ensured that cells which have a terminal missegregation at end of life do not show an inaccurately low histone level for the last cell cycle.

Following manual scoring of birth events, the fluorescent channel containing the histones was used to observe the missegregation dynamics. To ensure consistent scoring across experiments and eliminate bias, information about the experiments was masked from the scorer until after the data was evaluated. Following genome level missegregations, RETRN events were defined as where the histone fluorescence decreased in the daughter cell while simultaneously increasing in the mother cell. This prevented any changes in focus or gradual fluorescence increases due to fluorophore maturation from being inadvertently scored as a RETRN event. RETRN events were scored at the timepoint that the histones return to the mother cell. During cell cycles where cells had multiple retrograde events during the same cell cycle, only the final RETRN event was scored. Events were scored as terminal missegregation events if, prior to a correction, the daughter cell visibly separated from the mother cell (indicating cytokinesis) or if the mother died.

### Fluorescence quantification

Quantification of the level of protein localized to either the nucleolus (Cdc14) or the nucleus (Whi5), was done using a measure of how asymmetrically distributed the fluorescent signal was. Specifically, we used average brightness of the top 2% of pixels, divided by the cell median. By normalizing to median fluorescence, we corrected for any changes in fluorescence that could occur as a result of photobleaching. This method has been used previously as an accurate measure of the fraction of protein that is nuclear localized(Cai et al., 2008; Granados et al., 2017). For Myo1 quantification, we used the mean fluorescence level along the periphery of the cell as segmented. The nucleolar localized fluorescent protein (Utp13:GFP) was used to segment the nucleolus and determine the nucleolar size. This was done by calculating a segmentation threshold using Otsu’s threshold(Otsu, 1979) for each cell applied to the maximum projection of the fluorescent image stack. This approach was found to have good agreement with earlier estimates of nucleolar size in young cells grown in glucose(Jorgensen et al., 2007).

Because we are utilizing an epi-fluorescent microscope and as a result of out-of-focus light, mean fluorescence is not directly linked to protein concentration (Bakker, 2016). Because the fluorescence levels change dramatically during aging, segmenting out the nucleus to determine fluorescence levels could introduce significant biases as the cells age. Additionally, taking the mean fluorescence level could introduce significant errors depending on the segmentation accuracy. Instead, to normalize for changes, we sum a constant area within each cell. This prevents segmentation (of either the nucleus or cell) from contributing to the determination of histone amount. Pixels are sorted from high to low, and the top fraction corresponding to a circle with diameter of 3.8 μm are used to calculate the mean.

### Yeast Strains and Growth

The GFP strains were all acquired from the yeast GFP collection (Huh et al., 2003). The Htb2:mCherry strain was created by mating and sporulation of the strain from (Granados et al., 2017). This strain was then crossed with the relevant GFP strains (Nup49:GFP, Myo1:GFP, Tub1:GFP, Spc72:GFP, Cdc14:GFP, Utp13:GFP) or deletion strains (*hpc2*Δ, *fob1*Δ, *spt21*Δ, *tom1*Δ, *mad3*Δ) from the deletion collection (Winzeler et al., 1999) and then confirmed by colony PCR. The LacI-GFP strain with 50 LacO repeats on ChrXII was obtained from (Ide et al., 2010). The strains containing TetR-GFP and TetO repeats integrated into ChrIV or ChrV were obtained from (Fernius and Marston, 2009). These were then crossed with the strain containing Htb2:mCherry. Complete list of strains available in Table S1.

Prior to each experiment, single colonies were picked into SC media (Sunrise Biosciences) with 2% dextrose. Cells were grown overnight, and then diluted 1:200 in fresh media and grown for 5-6h. Prior to loading into the microfluidic device, 0.5 mL of SC 2% dextrose with 0.5% BSA was added to each 5 mL culture to prevent the cells from adhering to the PDMS during loading. During experiments, SC media with 2% dextrose and 0.1% BSA was used, and cells were imaged for 72h.

### Statistical Analysis

Error bars in the figures which contained bar plots were generated by bootstrapping with replacement, and then determining the 95% confidence intervals. Error bars in figures with line plots are standard error. Statistical significance for lifespan was determined using the log-rank test. Log-rank test was performed with, and without, censored cells that were lost prior to senescence or death. To compare distributions (such as numbers of missegregation events over the lifespan), a two-tailed t-test assuming equal variance was used. Correlations between histones or missegregation events and remaining replicative lifespan were calculated with the Spearman correlation using the population of cells alive at each replicative age. Correlation between Htb2:mCherry levels and DAPI staining were also done using the Spearman correlation.

## Supplementary Text

### Differences between censored and uncensored survival data

Frequently in experiments or clinical studies that involve the generation of survival curves, some samples will be removed from the population under observation. For example, a patient may leave a study not because of death, but because they move to a different country. This can be treated in a relatively straightforward manner statistically including these individuals in the analysis until the point that they are lost (or censored). This relies on the assumption that there is no bias in whether a sample is lost or retained. A recurring concern with microfluidic aging experiments involving yeast is whether there is a bias in how cells are lost or retained. This appears especially important when the mutation or transgene affects cell morphology or cell cycle, as this can result in a bias in which cells are lost from the traps. To reduce the likelihood that our observations were directly affected by loss rates, in the main text we have plotted all cells that were present at that replicative age for plots from birth. Thus, if a cell was lost at replicative age 20, it was included in the plots until age 19. In the supplementary materials we have also repeated every plot, but only using cells that died in the microfluidic device. Given that this is an altered population distribution and smaller number of cells, these plots are slightly different, but they do not affect the conclusions. Furthermore, in the plots where cells are aligned by birth rather than death, we are forced to only use cells that die in the device. For replicative lifespans shown in the supplementary, we include survival curves with and without censored cells.

### Aligning cells from birth or from death

Cells can be aligned either by birth (counting up from replicative age = 0), or by death (counting back from death). Either processing makes some assumptions about how similar cells are to one another. If cells are most similar to each other when they are born, aligning by birth makes sense, and as the replicative age increases, the number of samples decreases because cells are removed by death or senescence. In contrast, assuming that cells are similar at death implies that the phenotype of interest is most similar as cells approach death. For example, the average time cells take to proceed through each cell cycle increases geometrically when cells are aligned by birth, but exponentially when aligned by death. In the supplementary figures, we show the population means aligned using both approaches.

### Differences between Htb2:mCherry and Hta2:GFP

Although both of these strains were found to have similar numbers of missegregation events during their lifetimes, and similar fractions of these events were corrected (Figure 1 – figure supplement 4B,C), there are subtle differences between the strains. Most notably, the strain with Hta2:GFP had what we consider to be a normal lifespan for this background (median RLS= 21 for cells that died/senesced in the device, median RLS=24 including censored cells, Figure 1 – figure supplement 4D,E). The strain with Htb2:mCherry, however, had a somewhat shorter lifespan (median RLS = 16 for cells that died/senesced, median RLS=18 including censored cells, Figure 1 – figure supplement 4D,E). Removing FOB1, however, results in an increase of the replicative lifespan of this strain by ~30% (Figure 3 – figure supplement 3), which is in line with results from literature (McCormick et al., 2015). Furthermore, the increase in replicative lifespan as a result of increased histone transcription has been less thoroughly studied, but our results are in line with those previously reported by another group (Feser et al., 2010; Kruegel et al., 2011). Thus, although there is an unexpected reduction in lifespan for the Htb2:mCherry strain, we do not believe that it affects our results.

Likewise, as shown in the main text, we determined the correlation between missegregation events and remaining lifespan at the single cell level. The correlation is between the binary presence or absence of a missegregation event during a cell cycle and the remaining lifespan. Strikingly, as shown in Figure 1 – figure supplement 2, for both strains, the correlation between missegregation events and remaining replicative lifespan is the same for both Htb2:mCherry and Hta2:GFP. This is in spite of the difference in absolute lifespan between the two strains.

Because GFP fluorescence is much more affected than mCherry fluorescence by changing pH (Shaner et al., 2005), and the pH of the cytoplasm in aging yeast has previously been shown to increase (Henderson et al., 2014), we chose to perform the majority of the experiments using mCherry. This ensured that any changes in pH homeostasis during aging would not affect our measurements of histone levels.

### GFP Tagged Histones and Correlation with remaining RLS

As discussed in the main text, we imaged GFP tagged histones acquired from the GFP collection (Hta2, Htb2). This allowed us to determine that the increase in histone levels was a general aspect of aging physiology, not confined to specific histones. In order to determine whether histone levels were accurate biomarkers that predicted remaining lifespan, we used the single cell measurements collected for both strains (Figure 3 – figure supplement 1). At each replicative age, cells still alive are used to determine the correlation between their remaining replicative lifespan and the single cell histone levels. Thus, for prediction of remaining lifespan at replicative age 5, only cells that bud more than 5 times are included. The negative correlation between histone levels and remaining lifespan is similarly true for the GFP tagged histones imaged: Hta2, Htb2 (Figure 3 – figure supplement 1A). Furthermore, all histones are similarly predictive of remaining lifespan, and become increasingly predictive as cells age.

To provide a more complete picture of the data acquired, we plotted the GFP mean histone levels where single cells are aligned based on their replicative ages (Figure 3 – figure supplement 1B). As the population ages, the underlying distribution changes as cells die and are removed from the pool. This creates some variability with the increasing replicative age of the population. When individual cells are aligned by death, the population mean for histone levels at replicative ages preceding death looks quite different (Figure 3 – figure supplement 1C). By aligning cells by death, it is clear that histone levels begin to increase around 5-10 replications prior to death. When including all cells, including those that were lost from the traps, cells expressing GFP tagged histones have similar replicative lifespans (Figure 3 – figure supplement 1D,E).

### Single Cell DNA Levels and FOB1

As discussed in the main text, we determined the correlation between histone levels and DNA by comparing, in individual cells, levels of Htb2:mCherry with levels of DAPI. To do this, mother cells were aged in the device for 16h, and then fixed with ethanol and stained with DAPI. To determine whether the accumulation of ERCs with age leads to a detectable increase in DNA content, we compared DAPI staining of wild-type and *fob1*Δ. We acquired age-matched mother cells by using a multi-chamber device to simultaneously age wild-type and *fob1*Δ cells.

Comparing mothers that had grown in the device for 16h (median of 13 divisions and thus middle aged), we confirmed that DNA levels increase in wild-type cells are significantly higher compared with fob1Δ. The confirmed that the deletion of FOB1 decreased DNA content in aging cells, and not just histone levels. By manually scoring the number of divisions each strain had up until the point of ethanol fixation, we determined that there was no difference between strains in the number of daughters (Figure 3C, p=0.42). Although the replicative ages for each strain were statistically indistinguishable, the wild-type strain had a statistically significant increase in the DAPI staining levels (Figure 3D, p=0.0002). Thus, *fob1*Δ cells accumulate both histones and excess DNA at lower rates compared with wild-type cells. Furthermore, we compared Htb2 and DAPI levels at the single cell level, and histone levels are highly predictive of DNA content at the single cell level (Figure 1 – figure supplement 3).

### Manipulating histone demand (fob1Δ)

To provide a more complete picture of how Htb2 levels changed during aging in *fob1*Δ cells compared with wild-type, we performed the same procedure as with the GFP strains but using only the cells that die in the device to complement the plots in the main text. These plots look very similar to the plots using censored cells indicating that censoring doesn’t alter the population distribution. By aligning cells based on the replicative age, and plotting the mean Htb2:mCherry levels of the population, it is clear that there is a dramatic difference between wild-type and *fob1*Δ cells in Htb2 levels (Figure 3 – figure supplement 3A). This also reflects the missegregation rate (Figure 3 – figure supplement 3B). Missegregations are less predictive of death in *fob1*Δ cells than wild-type cells (Figure 3 – figure supplement 3C). When cells are aligned by death, as opposed to birth, both wild-type and fob1Δ cells experience an increase in histone levels, although *fob1*Δ cells are attenuated relative to wild-type (Figure 3 – figure supplement 3D). Because *fob1*Δ cells live longer, it would be reasonable to expect that by the end-of-life, the missegregation rate is closer between wild-type and *fob1*Δ cells than when aligned by birth (Figure 3 – figure supplement 3E). Similar to previous reports (Defossez et al., 1999), we find a significant increase the replicative lifespan of *fob1*Δ cells compared with wild-type (Figure 3 – figure supplement 3F). Survival curves of only the cells which died in the device, and thus were used in the analysis, showed similar lifespan differences (Figure 3 – figure supplement 3G).

### Manipulating histone supply (*tom1*Δ, *hpc2*Δ and *spt21*Δ)

To provide a more complete picture of how the manipulation of the histone supply affected lifespan and Htb2 levels, we performed a similar procedure as previously described. Spt21 positively regulates the transcription of all histone genes, and Hpc2 negatively regulates histone genes. The protein Tom1 is involved in the ubiquitination and degradation of excess histone proteins. Thus by deleting *HPC2*, we upregulate histone transcription, deleting *SPT21* suppresses histone transcription and deleting *TOM1* allows excess histone proteins to remain in the cell. Similar to the figures in the main text, we used only cells that die in the device to plot missegregation rates aligned by birth (Figure 5 – figure supplement 1A) and by death (Figure 5 – figure supplement 1B). As previously identified, *hpc2*Δ and *tom1*Δ cells live significantly longer than wild-type cells (Figure 5 – figure supplement 1C,D). Surprisingly, *spt21*Δ cells do not appear shorter lived compared to wild-type (Figure 5 – figure supplement 1C,D). These trends are maintained in survival curves of only cells that die in the device (Figure 5 – figure supplement 1C).

**Table S1.** List of all strains and genotypes used in this study.

| Strain Name | Genotype | Source |
| --- | --- | --- |
| BY4741 | MATa his3Δ1 leu2Δ0 met15Δ0 ura3Δ0 |  |
| BY4742 | MATα his3Δ1 leu2Δ0 lys2Δ0 ura3Δ0 |  |
| YSI129 | ade2-1 ura3-1 his3-11 trp1-1 leu2-3,112 can1-100 fob1Δ::LEU2 his3-11::GFP-LacI-HIS3 LacO(50)-ADE2:445kb ChXII (110 rDNA copies) | Ide, et al. 2010 |
| AMY914 | MATa ura3 trp1 his3-11 MET-CDC20::URA pURA::tetR-GFP::LEU2 cenIV::tetOx448::URA3 | Fernius, et al. 2009 |
| AMY1081 | MATa ade2-1 leu2-3 trp1-1 trp1 his3-11 can1-100 GAL psi+ MET-CDC20::URA pURA::tetR-GFP::LEU2 ura3::tetOx112::URA3 | Fernius, et al. 2009 |
| MC213 | HTB2::mCherry-URA3 his3 leu2 met15 LYS2 | this study |
| MC230 | fob1Δ::KanMX his3 leu2 HTB2::mCherry-URA met15 LYS2 | this study |
| MC237 | HTA2::GFP-HIS3 his3 leu2 ura3 met15 LYS2 | Yeast GFP Collection |
| MC239 | HTB2::GFP-HIS3 his3 leu2 ura3 met15 LYS2 | Yeast GFP Collection |
| MC245 | NUP49::GFP-HIS3 his3 leu2 ura3 met15 LYS2 | Yeast GFP Collection |
| MC247 | TUB1::GFP-HIS3 his3 leu2 ura3 met15 LYS2 | Yeast GFP Collection |
| MC250 | HTB2::mCherry-URA3 WHI5::GFP-HIS3 leu2 met15 LYS2 | this study |
| MC255 | spt21Δ::KanMX his3 leu2 HTB2::mCherry-URA met15 LYS2 | Yeast KO Collection |
| MC257 | hpc2Δ::KanMX his3 leu2 HTB2::mCherry-URA met15 LYS2 | Yeast KO Collection |
| MC258 | NUP49::GFP-HIS3 his3 leu2 ura3 met15 LYS2 htb2::mCherry-URA | this study |
| MC263 | HTB2::mCherry-URA3 his3 leu2 met15 lys2 hpc2Δ::KanMX | this study |
| MC264 | HTB2::mCherry-URA3 TUB1::GFP-HIS2 leu2 met15 LYS2 | this study |
| MC266 | HTB2::mCherry-URA3 his2 leu2 MET15 lys2 spt21Δ::KanMX | this study |
| MC281 | HTB2::mCherry-URA3 his3-11::GFP-LacI-HIS3 RDN1::LacO(50)-ADE2 | this study |
| MC349 | mad3Δ::KanMX his3 leu2 htb2::mCherry-URA met15 LYS2 | this study |
| MC351 | SPC72::GFP-HIS3 his3 leu2 ura3 met15 LYS2 | Yeast GFP Collection |
| MC352 | UTP13::GFP-HIS3 his3 leu2 ura3 met15 LYS2 | Yeast GFP Collection |
| MC354 | CDC14::GFP-HIS3 his3 leu2 ura3 met15 LYS2 | Yeast GFP Collection |
| MC355 | SPC72::GFP-HIS3 his3 leu2 ura3 met15 LYS2 HTB2:mCherry-URA | this study |
| MC360 | CDC14::GFP-HIS3 his3 leu2 ura3 met15 LYS2 HTB2:mCherry-URA | this study |
| MC364 | UTP13::GFP-HIS3 his3 leu2 ura3 met15 LYS2 HTB2:mCherry-URA | this study |
| MC367 | mad3Δ::KanMX his3 leu2 ura3 met15 LYS2 HTB2:mCherry-URA | this study |
| MC372 | tom1Δ::KanMX his3 leu2 ura3 met15 LYS2 HTB2:mCherry-URA | this study |
| MC394 | ura3 his3-11 pURA::tetR-GFP::LEU2 cenIV::tetOx448::URA3 HTB2:mCherry-URA | this study |
| MC395 | leu2-3 his3-11 pURA::tetR-GFP::LEU2 ura3::tetOx112::URA3 HTB2:mCherry-URA | this study |

**Figure 1 – figure supplement 1.**
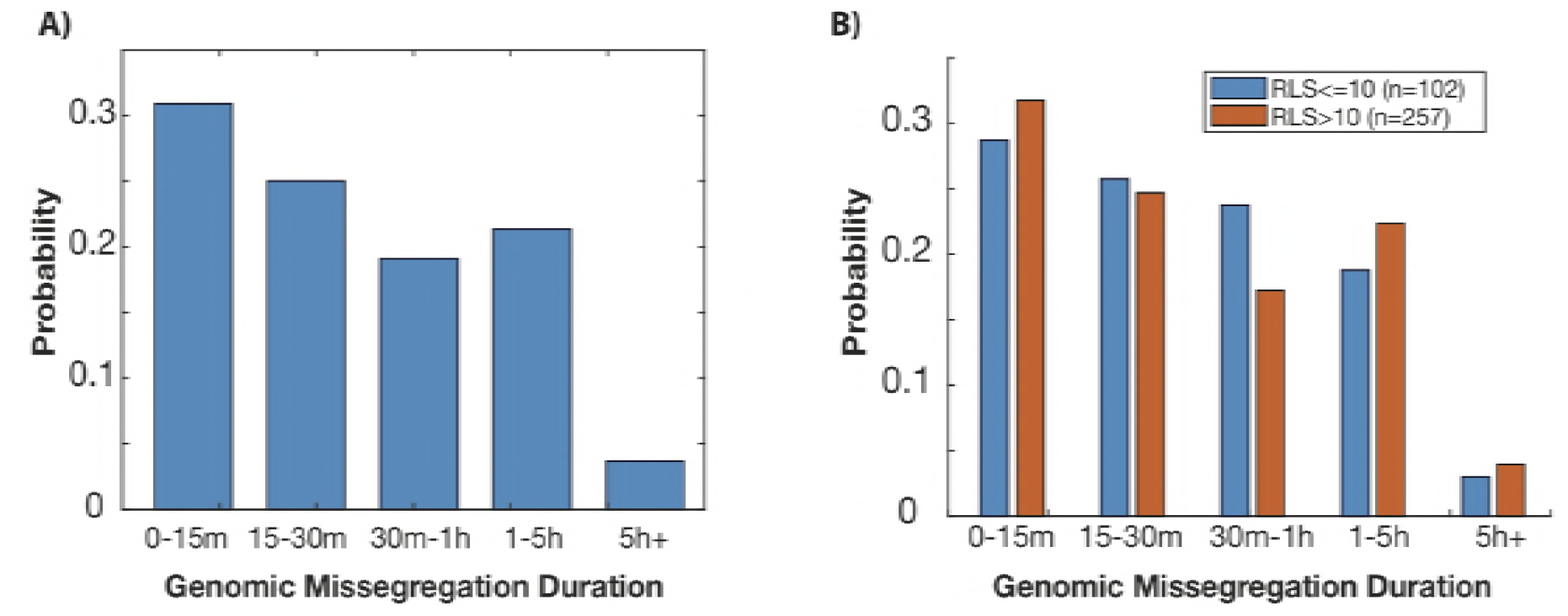
Histogram showing the duration of genomic missegregation events in wild-type cells. Only events which eventually resulted in a RETRN event are shown, as that provides a clear end of the missegregation event. **A**) Many GLMs are corrected within an hour, but some events can last several hours. This includes GLMs from cells of all replicative ages. **B**) When events are separated based on the age of the mother cell at time of missegregation event, there is no significant difference in the distribution of GLM durations (p=0.5, two-tailed t-test). N-value are the number of individual missegregation events quantified.

**Figure 1 – figure supplement 2.**
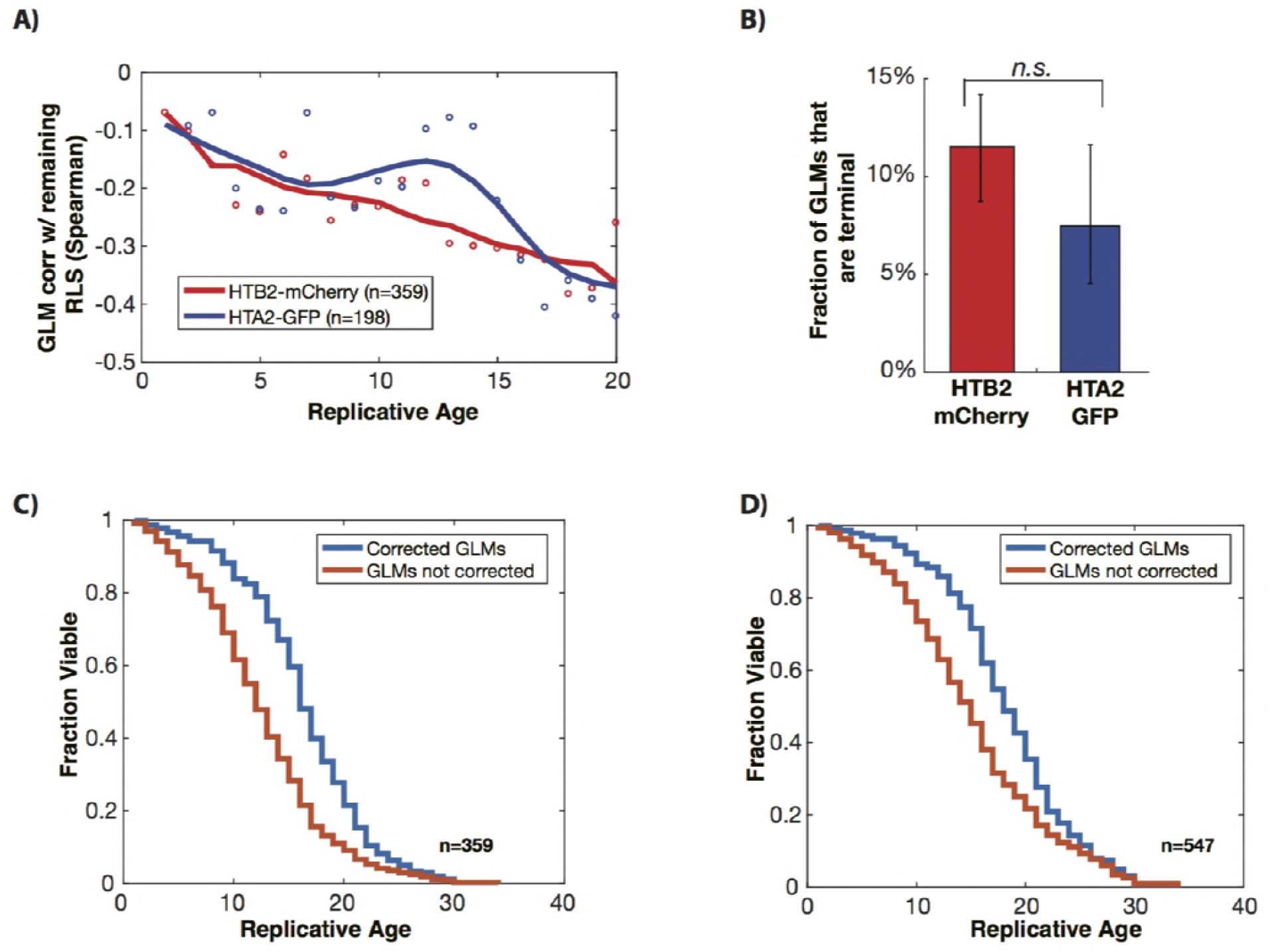
GLMs are predictive of death, and most GLMs are corrected by RETRN events. **A**) At the single cell level, genome level missegregation (GLM) events are correlated with impending mortality (anti-correlated with remaining lifespan, as shown), and become more predictive (more strongly anti-correlated, as shown) with increasing replicative age. The strength of the anti-correlation with age is similar regardless of the fluorophore used or histone protein tagged. Dots show correlation between an individual histone and the remaining lifespan at that replicative age. To show trends, these have been smoothed with a moving average (solid line). **B**) When a GLM occurs, approximately 90% of the time it is corrected through a RETRN event, and the remainder of the time it becomes a terminal missegregation. Error bars are 95% confidence intervals generated by bootstrapping with replacement. N-values were individual cells, and only cells that died or senesced in the device were used for both plots. **C**) RETRN events significantly increase replicative life span. Comparison of experimentally observed replicative lifespan (corrected GLMs) to expected lifespan if all GLMs were terminal (not corrected) for wild type cells. To generate the GLM not corrected lifespan, the first GLM event for each mother cell was assumed to be terminal, and the lifespan was truncated accordingly. This plot uses only cells that die in the device. RETRN events increase median lifespan by 33% (p<0.0001). **D**) Same is in panel S2C, but including cells that are censored. The increase in median lifespan by RETRN events is 29% (p<0.0001).

**Figure 1 – figure supplement 3.**
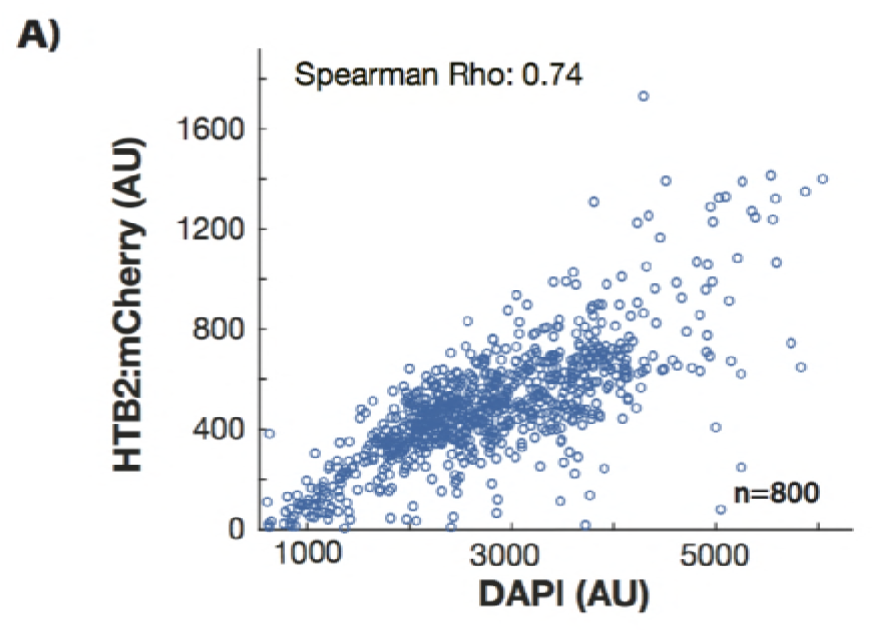
Htb2 levels are predictive of DNA content at the single cell level. Wild-type cells were grown in the microfluidic device for 16h, and then fixed. Cells had a median replicative age of 13 divisions. The level of Htb2:mCherry is highly correlated (p<0.0001) with the level of DAPI staining at the single cell level in individual cells. This is true across the entire range of Htb2:mCherry fluorescence, as even cells that have the highest Htb2:mCherry levels have correspondingly high DAPI staining. N=800 individual mother cells.

**Figure 1 – figure supplement 4.**
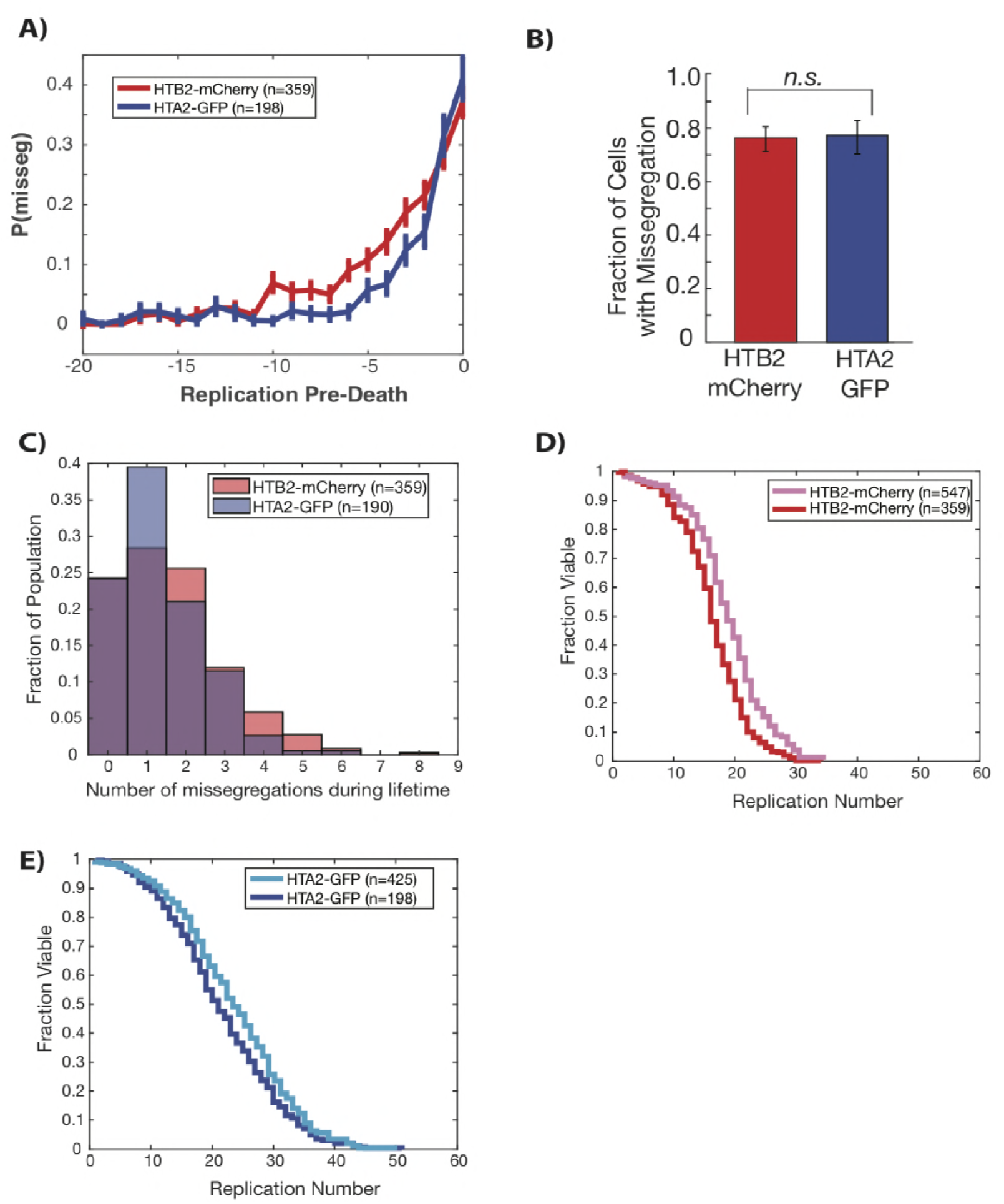
GLMs increase at end of life regardless of histone tagged and fluorophore used. **A**) Both Htb2:mCherry and Hta2:GFP strains experience a dramatic increase in the probability of GLM events in the 5-6 divisions preceding death. Error bars are standard error. **B**) There is no difference between Htb2:mCherry and Hta2:GFP with respect to the fraction of cells that experience GLM events. Error bars are 95% confidence intervals from bootstrapping over individual cells. **C**) The distribution of the number of GLM events that cells experience over their lifetime is similar between the strains but statistically different (p=0.02 two-tailed t-test), with Htb2:mCherry cells experiencing, on average, more events over their lifetime. **D**) Replicative lifespan curves for Htb2:mCherry including censored cells (pink) and excluding (red). **E**) Replicative lifespan curves for Hta2:GFP including censored cells (light blue) and excluding (blue).

**Figure 1 – figure supplement 5.**
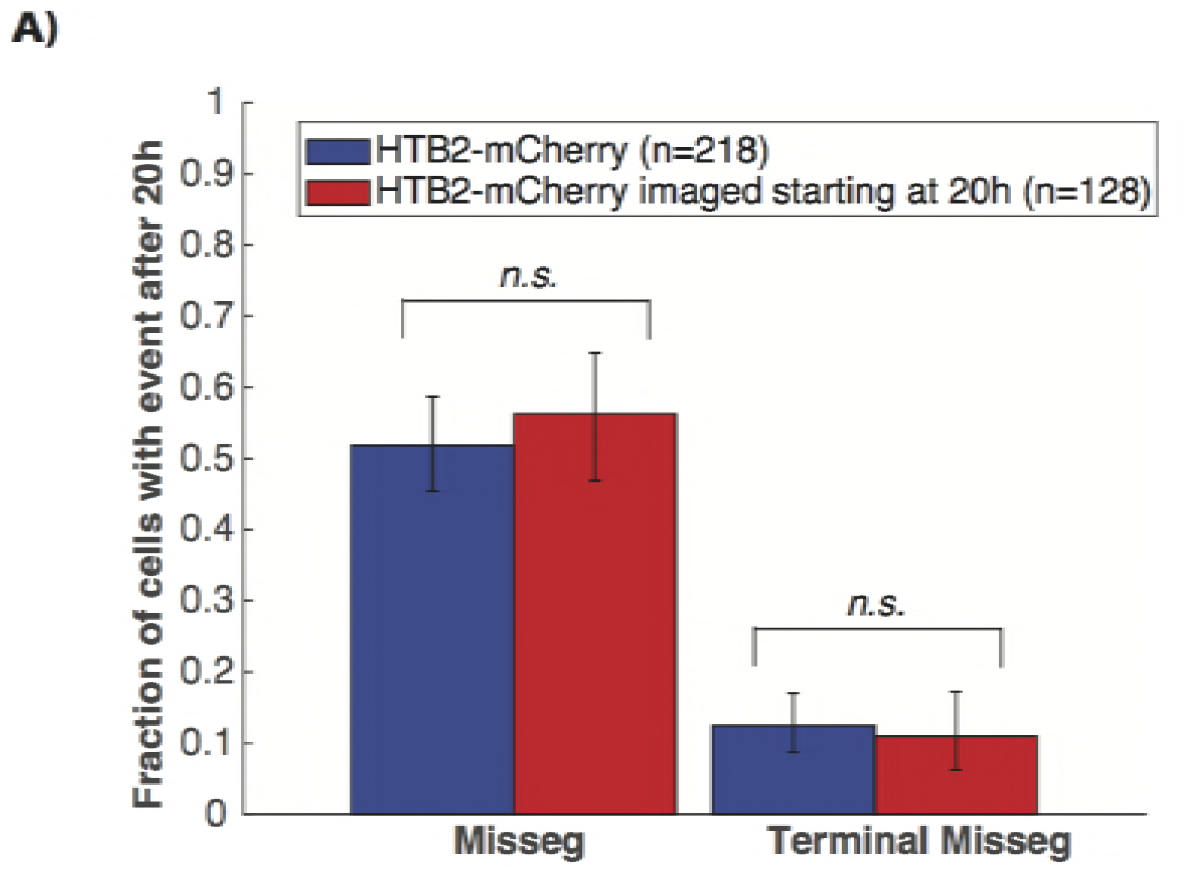
GLMs are not caused by imaging conditions. **A**) To determine whether cells were affected by the cumulative exposure to fluorescence excitation energy, we compared GLM rates in cells imaged over their entire lifespans (blue) with those only imaged after they were already aged for 20 hours (red), corresponding to a median replicative age of 16 generations. This is equivalent to approximately 75% of the median lifespan of this strain. To compare these cells to the control, we only quantified GLMs that occurred after 20h. Error bars are 95% confidence intervals generated by bootstrapping with replacement over all cells. No difference was detected in cells imaged continuously over their entire lifespan or only after 20 hours, indicating that there is no cumulative effect of the exposure to fluorescence excitation light on the frequency of GLMs or the ability of cells to correct these through RETRN events.

**Figure 2 – figure supplement 1.**
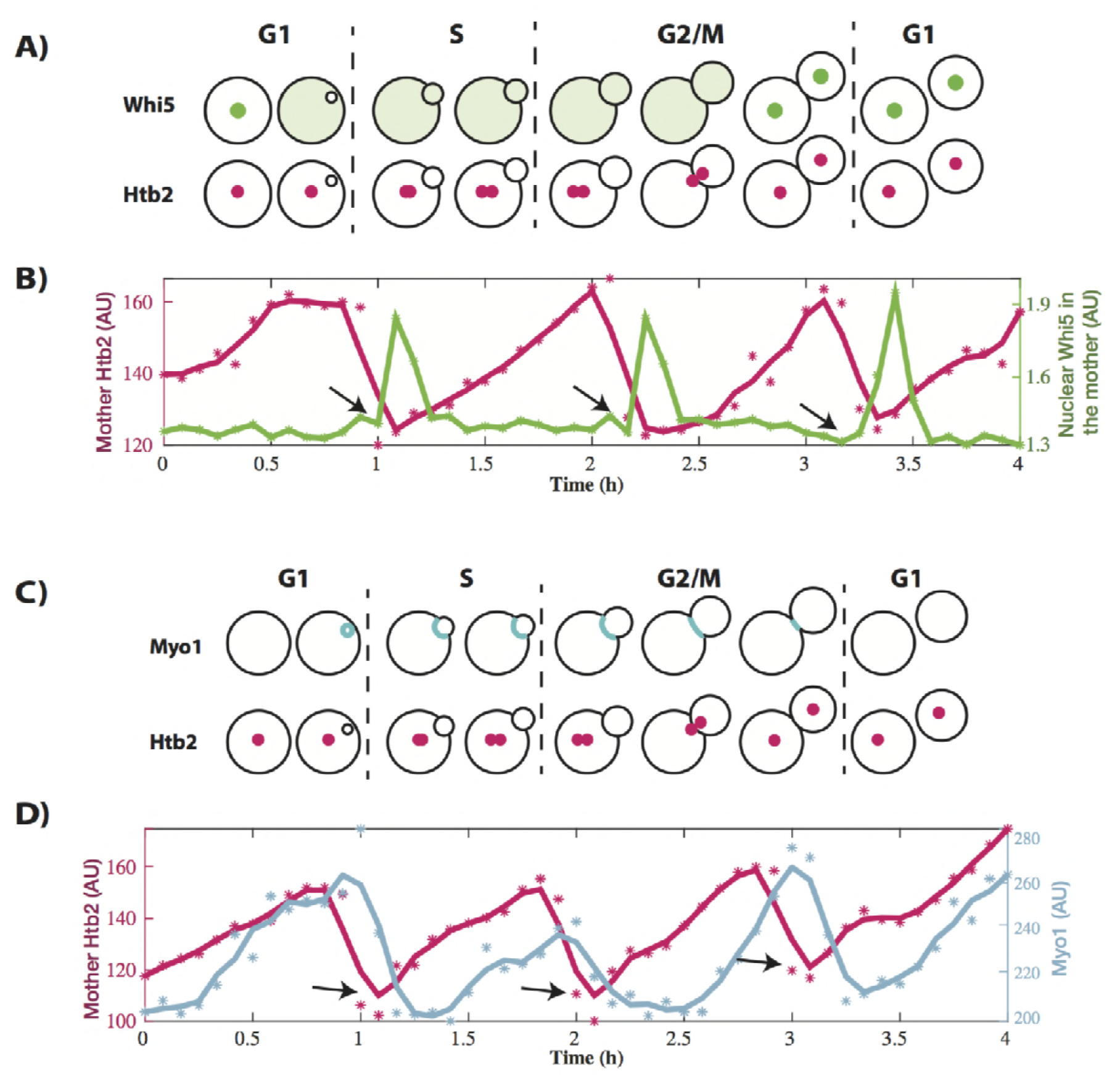
Single cell traces of the Whi5 and Myo1 data. **A**) Schematic showing the temporal dynamics of the proteins used to characterize missegregation events. Whi5 exits the nucleus to initiate START and move the cell into S phase. It re-enters the nucleus at the end of mitosis. **B**) A representative trace of a single cell expressing Htb2:mCherry and Whi5:GFP. Histone levels (pink) increase, and then fall during mitosis. Arrows indicate the timepoint before Whi5 transitions from the cytoplasm to the nucleus. (*) indicate raw Htb2 measurements, which were smoothed using a moving average for legibility (solid line). **C**) Schematic showing the temporal dynamics of Myo1. Myo1 is produced at the end of G1 and localizes to the bud neck until cytokinesis ends mitosis. **D**) Representative single cell trace of a cell containing Htb2:mCherry and Myo1:GFP. (*) indicate raw Htb2 or Myo1 measurements, which were smoothed using a moving average for legibility (solid line). Arrows indicate the lowest point of Htb2 levels following mitosis.

**Figure 2 – figure supplement 2.**
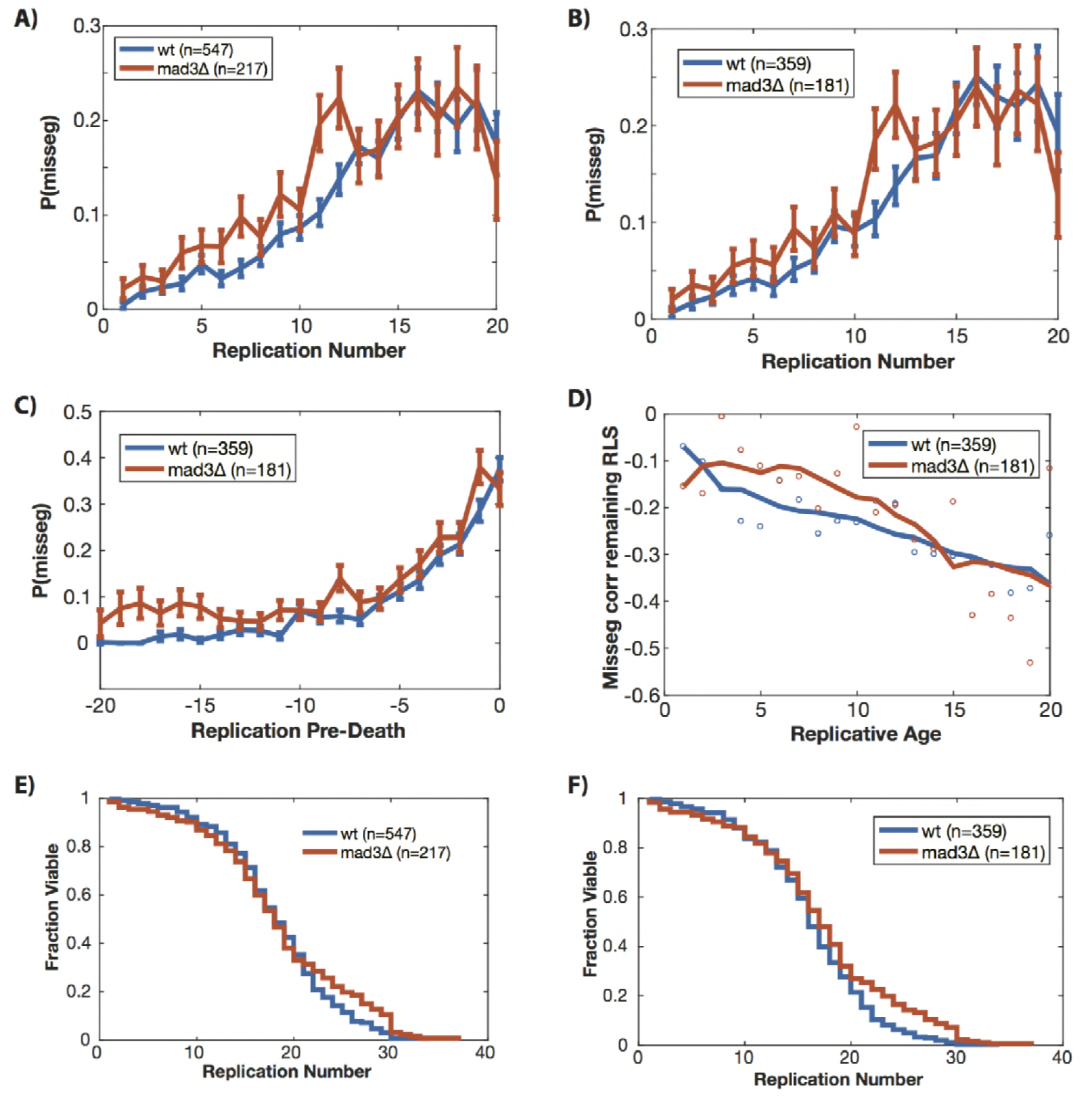
Removing Mad3 (mammalian BubR1) fails to affect the age-related increase in missegregation rate. **A**) Wild-type and *mad3*Δ cells experience a similar increase in genome level missegregation (GLM) events with age when looking at all cells. **B**) The increase in GLM rate is similar when only comparing cells that die or senesce in the device. **C**) When only cells that die in the device are aligned by death, both *mad3*Δ and wild-type cells experience a similar age-related increase in GLM rates that begins around 5 divisions prior to death. **D**) At the single cell level, GLM events are equally strongly correlated with impending mortality in both *mad3*Δ and wild-type cells. Dots show correlation between a GLM and the remaining lifespan at that replicative age for that genotype. To show trends, these have been smoothed with a moving average for legibility (solid line). **E**) RLS curve including censored cells (used in A). **F**) RLS curve excluding censored cells (used in B-D). All error bars are standard error.

**Figure 3 – figure supplement 1.**
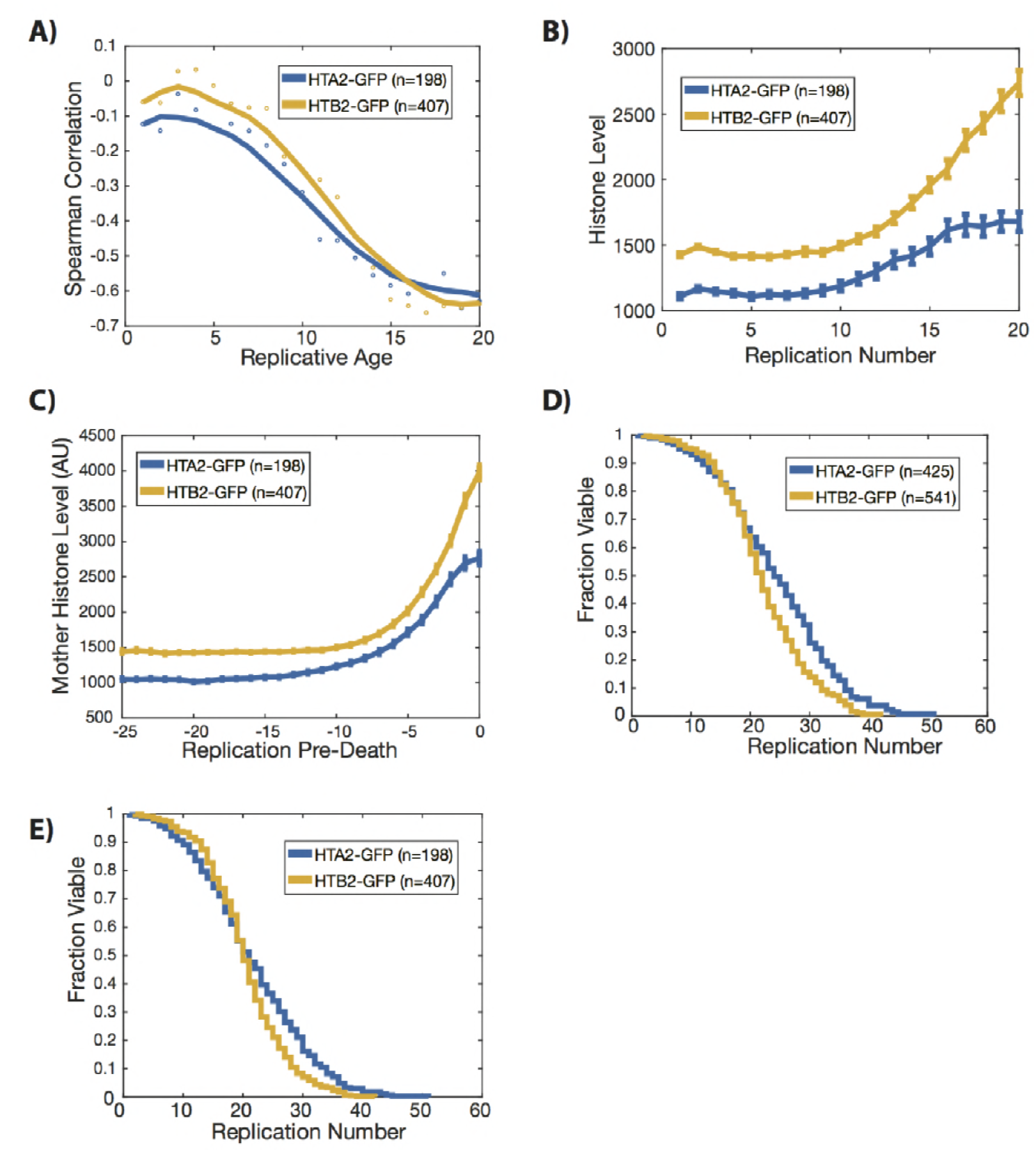
GFP tagged histone data. Histones tagged with GFP increase during aging, and are predictive of mortality. **A**) At the single cell level, histone levels are strongly correlated with impending mortality, regardless of the histone tagged. Histones become increasingly negatively correlated with remaining replicative lifespan as the cells age. Dots show correlation between an individual histone and the remaining lifespan at that replicative age. To show trends, these have been smoothed with a moving average (solid line). **B**) Mean histone levels at each division aligned based on replicative age for each cell. **C**) Mean histone levels at each division aligned using individual cell death. This shows that, on average, all histones begin to increase sometime around 5-10 divisions prior to death. **D**) The replicative lifespans of the strains including censored cells. **E**) The replicative lifespan of only cells that die or senesce in the device. These are the cells that are used in the analysis for A-C. N values indicate the number of cells, and the error bars are standard error.

**Figure 3 – figure supplement 3.**
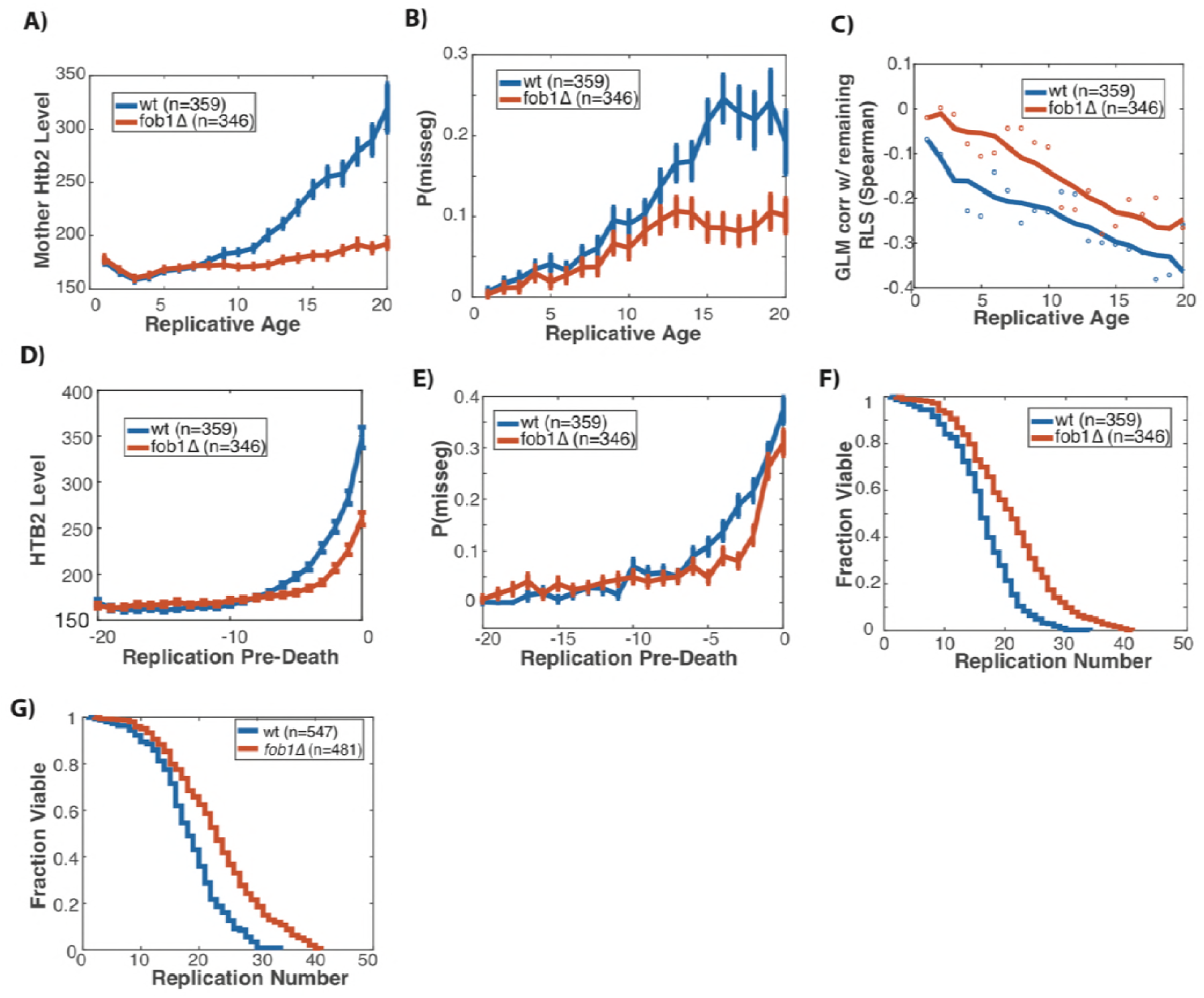
Fob1 vs Wild Type data. **A**) Mean Htb2 levels of mother cells aligned by birth and excluding cells that were censored (See Figure 3B) **B**) Mean missegregation rates of wild-type and *fob1*Δ mother cells aligned by birth and excluding cells that were censored (See Figure 4A). **C**) At the single cell level, missegregation events are less strongly anti-correlated with remaining lifespan in *fob1*Δ compared with wild-type. The correlation curve for*fob1*Δ cells appears parallel to wild-type, but with an offset that is roughly proportional to the increase in replicative lifespan we observed for the *fob1*Δ cells. Excludes censored cells. Dots show correlation between an individual histone and the remaining lifespan at that replicative age. To show trends, these have been smoothed with a moving average (solid line). **D**) Mean Htb2 levels of mother cells aligned by death instead of birth, excluding censored cells. **E**) Probability of missegregation events for wild-type and *fob1*Δ cells, but aligned by death instead of birth and excluding censored cells. **F**) Survival curve of wild-type and *fob1*Δ cells containing only cells which are not lost and censored. These are the cells used in A-E. Survival curves comparing wild-type cells and cells lacking *FOB1*. **G**) Survival curve including censored cells lost during the experiment. Compared with wild-type cells, *fob1*Δ mutants live longer (p<1E-10, log-rank test). These are the cells used in the main figures. N-values report the number of cells, and error bars for C,D,F are standard error.

**Figure 4 – figure supplement 1.**
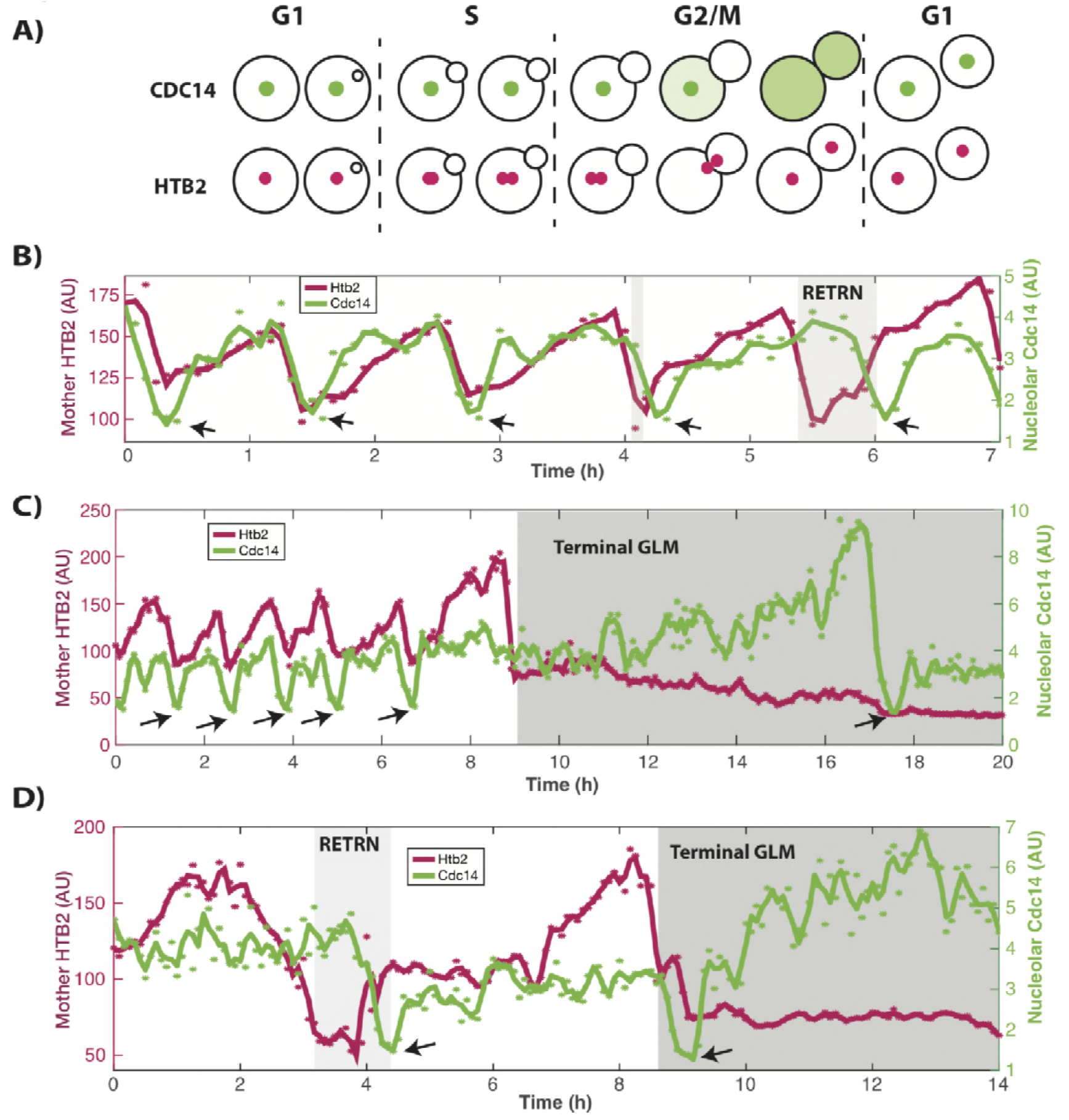
Cdc14 single cell traces in cells with terminal GLM and RETRN events. **A**) Schematic showing cell cycle dynamics. Cdc14 is localized to the nucleolus during the majority of the cell cycle and exits it in two stages during mitosis. Cdc14 re-enters the nucleolus at the end of mitosis. **B**) A representative trace of a single cell expressing Htb2:mCherry and Cdc14:GFP showing normal divisions and RETRN events. Histone levels (pink) increase, and then fall during mitosis. Arrows indicate the timepoints where Cdc14 is at a minima in the nucleolus. In normal divisions this coincides with the Htb2 minima. (*) indicate raw Htb2 and Cdc14 measurements, which were smoothed using a moving average and Savitzky-Golay smoothing respectively for legibility (solid line). In the last two divisions, a RETRN events can be observed. **C**) Representative trace of a single cell expressing Htb2:mCherry and Cdc14:GFP showing a terminal missegregations. Cdc14 exits the nucleolus nearly eight hours after the GLM, but there is no RETRN event. **D**) Representative trace of a single cell expressing Htb2:mCherry and Cdc14:GFP showing a GLM with a RETRN event (4h) and then a terminal missegregation.

**Figure 5 – figure supplement 1.**
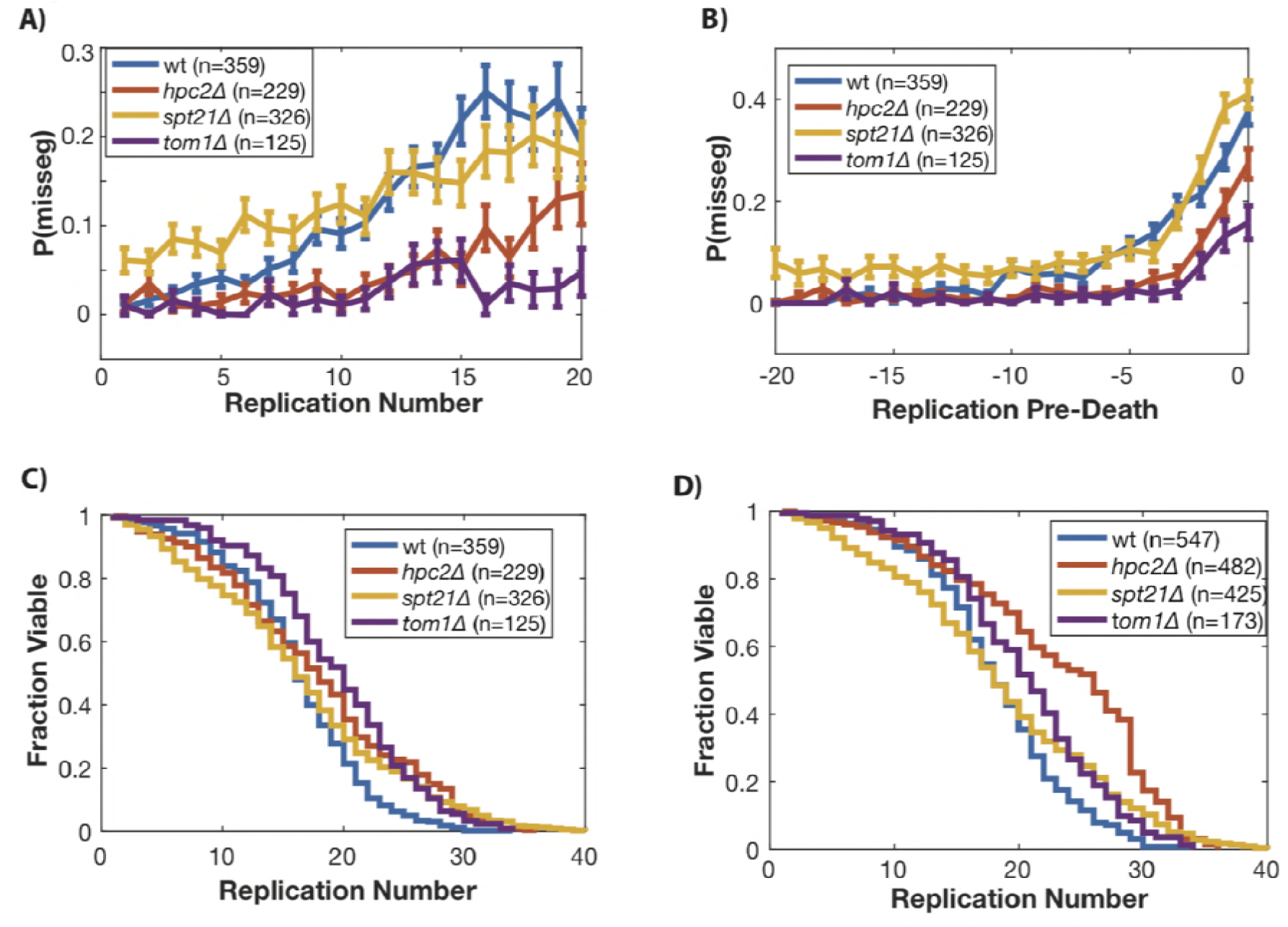
Manipulating histone supply in yeast affects genome level missegregation (GLM) rates. **A**) By manipulating the supply of histones through deleting *SPT21, TOM1* and *HPC2* we can affect the probability of GLMs occurring. **B**) Mean GLM rates for wild-type (wt), *spt21*Δ, *hpc2*Δ and *tom1*Δ mother cells aligned by birth and excluding cells that were censored (See Figure 4B). **C**) Survival curves of only cells that are not lost. These are the cells used for A,B. D) Survival curves which account for cells which are censored based on loss from the traps, for wt, *hpc2*Δ and *spt21*Δ and *tom1*Δ. These are the cells used in Figure 5 of the main text. Wild-type and *hpc2*Δ have different lifespans (p<0.001, log-rank test), wild-type and *tom1*Δ have different lifespans (p<0.001, log-rank test), and wild-type and *spt21*Δ do not have different lifespans (p=0.1 log-rank test). N-values indicate cell numbers, and error bars for A,B are standard error.

## Video Legends

**Video 1** – Normal divisions followed by RETRN events in a strain expressing Htb2:mCherry and Myo1:GFP. The mother cell undergoes four normal divisions, and on the fifth (at timepoint 7h:35min), it experiences a missegregation event. The bud neck is clearly maintained until the retrograde transport occurs at 12h. Following this event, the bud neck is quickly removed, and is completely gone by 12h:15min. The blue arrow points to the mother cell during timepoints where it is experiencing a missegregation event. Timestamp is Hours:Min

**Video 2** – RETRN events in a strain expressing Htb2:mCherry and Myo1:GFP. At timepoint 1h:20min the mother cell experiences a missegregation event. The bud neck is clearly maintained until the retrograde transport occurs at 2h:40min. Following this event, the bud neck is quickly removed, and is completely gone by 2h:55min. The blue arrow points to the mother cell during timepoints where it is experiencing a missegregation event. Video is significantly slower than other Videos. Timestamp is Hours:Min.

**Video 3** - DNA colocalizes with tagged histones through mitosis. Cells expressing Htb2:mCherry were stained with Hoechst 3342, a live DNA stain. The first part of the video shows an overlay of red (Htb2:mCherry) and blue (Hoechst 3342), and the second shows the channels separated. As can be clearly seen in a normal cell cycle, the histones colocalize with the DNA, and both increase or decrease in fluorescence in the mother cell simultaneously.

**Video 4** - DNA colocalizes with tagged histones through mitosis, including during missegregation and RETRN events. Cells expressing Htb2:mCherry were stained with Hoechst 3342, a live DNA stain. The first part of the video shows an overlay of red (Htb2:mCherry) and blue (Hoechst 3342), and the second shows the channels separated. As can be clearly seen, the histones colocalize with the DNA, and both increase or decrease in fluorescence in the mother cell simultaneously.

**Video 5** – Normal divisions in a strain expressing Htb2:mCherry and Nup49:GFP. This cell undergoes six divisions, with histone and nuclear envelope behavior that is characteristic of young, healthy cells. Timestamp is Hours:Min

**Video 6** – A missegregation followed by RETRN event in a strain expressing Htb2:mCherry and Nup49:GFP. The initial missegregation can be seen at timepoint 3h:30min, and the RETRN event at 8h:30min. Following the RETRN event, the mother cell is able to bud again at 12h, but the nuclear morphology of the daughter (for example, at 16h) is significantly altered. The blue arrow points to the mother cell during timepoints where it is experiencing a missegregation event. Timestamp is Hours:Min

**Video 7** – Terminal missegregations in a strain expressing Htb2:mCherry and Nup49:GFP. At 3h, the mother cell can be seen to undergo a missegregation event. At 15h:30min, the daughter cell buds and can be seen to undergo mitosis, indicating that the daughter cell has separated from the mother. The blue arrow points to the mother cell during timepoints where it is experiencing a missegregation event. The mother cell eventually dies at 40h. Timestamp is Hours:Min.

**Video 8** - A normal cell division, followed by a missegregation and RETRN event in a cell expressing Htb2:mCherry and Spc72:GFP. The two green dots indicate the spindle poles, and at numerous timepoints both poles enter the daughter cell. The blue arrow points to the mother cell during timepoints where it is experiencing a missegregation event. Timestamp is Hours:Min

**Video 9** - A terminal missegregation event in a cell expressing Htb2:mCherry and Spc72:GFP. The two green dots indicate the spindle poles, and both poles enter the daughter around 2h40m. The poles move around and are highly active, with one at times reentering the mother cell. Finally, at 5h:20m, the daughter cell is washed away indicating it has fully separated from the mother and that this is a terminal missegregation. The blue arrow points to the mother cell during timepoints where it is experiencing a missegregation event. Timestamp is Hours:Min.

**Video 10** – Normal divisions followed RETRN events in a strain expressing Htb2:mCherry and Tub1:GFP. The mother cell undergoes four divisions normally and on the fifth, at timepoint 7h:50min it experiences a missegregation event that is resolved by a RETRN event at 10h:30min. The blue arrow points to the mother cell during timepoints where it is experiencing a missegregation event. Timestamp is Hours:Min

**Video 11** – A terminal divisions followed RETRN events in a strain expressing Htb2:mCherry and Tub1:GFP. At timepoint 3h:15min the mother experiences a missegregation event, and both the chromatin and microtubules can be seen entering the daughter cell. At 7h:25 min the daughter cell is washed away indicating the cell has completed cytokinesis. The blue arrow points to the mother cell during timepoints where it is experiencing a missegregation event. Timestamp is Hours:Min

**Video 12** – Normal divisions, and an increase in histone levels in a strain expressing Htb2:mCherry and Nup49:GFP. Over a period of 15h this cell grows normally and produces daughters. The increase in Htb2:mCherry can be clearly seen over the course of this experiment.

**Video 13** - Age related structural and morphological changes to the nucleolus. This strain is expressing a nucleolar localized protein (Utp13:GFP) and Htb2:mCherry. In the young cell, the green nucleolus is adjacent to the red genomic material. As the cell ages and accumulates ERCs, the histone level begins to increase noticeably around 14h. This alters the nucleolar morphology and size, and by 23h the nucleolar morphology is dramatically different and it surrounds a large portion of the histones. Timestamp is Hours:Min.

**Video 14** - Age related structural and morphological changes to the nucleolus. This strain is expressing a nucleolar localized protein (Utp13:GFP) and Htb2:mCherry. In the young cell, the green nucleolus is adjacent to the red genomic material. As the cell ages and accumulates ERCs, the histone level begins to increase noticeably around 9h. This alters the nucleolar morphology and size, and by 12h the nucleolar morphology is dramatically different and it surrounds a large portion of the histones. Timestamp is Hours:Min.

**Video 15** - A missegregation and RETRN event showing that the Cdc14 remains localized to the nucleolus even when the cell experiences a missegregation event. The cell is expressing Htb2:mCherry and Cdc14:GFP. The exit of Cdc14:GFP from the nucleolus at 2h coincides with the RETRN event. The blue arrow points to the mother cell during timepoints where it is experiencing a missegregation event. Timestamp is Hours:Min

**Video 16** - ChrXII dynamics during a missegregation and RETRN event. Cell is expressing Htb2:mCherry and LacI:GFP, and has LacO repeats inserted into ChrXII. The Chromosome XII sister chromatids clearly remain behind despite the majority of the genome entering the daughter cell. Simultaneous to the RETRN event, the sister chromatids separate and can be identified in both mother and daughter cells. The blue arrow points to the mother cell during timepoints where it is experiencing a missegregation event. Timestamp is Hours:Min

**Video 17** - ChrXII dynamics during a missegregation and RETRN event. Cell is expressing Htb2:mCherry and LacI:GFP, and has LacO repeats inserted into ChrXII. The Chromosome XII sister chromatids clearly remain behind despite the majority of the genome entering the daughter cell. Simultaneous to the RETRN event, the sister chromatids separate and can be identified in both mother and daughter cells. The blue arrow points to the mother cell during timepoints where it is experiencing a missegregation event. Timestamp is Hours:Min

**Video 18** - ChrIV dynamics during a missegregation and RETRN event. Cell is expressing Htb2:mCherry and TetR:GFP, and has TetO repeats inserted into ChrIV. The Chromosome IV sister chromatids clearly move into the daughter with the majority of the genome. Simultaneous to the RETRN event, the sister chromatids separate and can be identified in both mother and daughter cells. The blue arrow points to the mother cell during timepoints where it is experiencing a missegregation event. Timestamp is Hours:Min

**Video 19** - ChrV dynamics during a missegregation and RETRN event. Cell is expressing Htb2:mCherry and TetR:GFP, and has TetO repeats inserted into ChrV. The Chromosome V sister chromatids clearly move into the daughter with the majority of the genome. Simultaneous to the RETRN event, the sister chromatids separate and can be identified in both mother and daughter cells. The blue arrow points to the mother cell during timepoints where it is experiencing a missegregation event. Timestamp is Hours:Min

**Video 20**- Cdc14 dynamics during normal cell divisions. Cell is expressing Htb2:mCherry and Cdc14:GFP. Timestamp is Hours:Min

**Video 21** - Spindle pole dynamics during normal cell divisions. Cell is expressing Htb2:mCherry and Spc72:GFP. Timestamp is Hours:Min

**Video 22** - Microtubule dynamics during normal cell divisions. Cell is expressing Htb2:mCherry and Tub1:GFP. Timestamp is Hours:Min

**Video 23**- Bud neck dynamics during normal cell divisions. Cell is expressing Htb2:mCherry and Myo1:GFP. Timestamp is Hours:Min

